# Comparative analysis of 43 distinct RNA modifications by nanopore tRNA sequencing

**DOI:** 10.1101/2024.07.23.604651

**Authors:** Laura K. White, Kezia Dobson, Samantha del Pozo, Jill M. Bilodeaux, Shelby E. Andersen, Amber Baldwin, Chloe Barrington, Nadine Körtel, Federico Martinez-Seidel, Saylor M. Strugar, Kristin E.N. Watt, Neelanjan Mukherjee, Jay R. Hesselberth

## Abstract

Transfer RNAs are the fundamental adapter molecules of protein synthesis and the most abundant and heterogeneous class of noncoding RNA molecules in cells. The study of tRNA repertoires remains challenging, complicated by the presence of dozens of post transcriptional modifications. Nanopore sequencing is an emerging technology with promise for both tRNA sequencing and the detection of RNA modifications; however, such studies have been limited by the throughput and accuracy of direct RNA sequencing methods. Moreover, detection of the complete set of tRNA modifications by nanopore sequencing remains challenging. Here we show that recent updates to nanopore direct RNA sequencing chemistry (RNA004) combined with our own optimizations to tRNA sequencing protocols and analysis workflows enable high throughput coverage of tRNA molecules and characterization of nanopore signals produced by 43 distinct RNA modifications. We share best practices and protocols for nanopore sequencing of tRNA and further report successful detection of low abundance mitochondrial and viral tRNAs, providing proof of concept for use of nanopore sequencing to study tRNA populations in the context of infection and organelle biology. This work provides a roadmap to guide future efforts towards *de novo* detection of RNA modifications across multiple organisms using nanopore sequencing.

## Introduction

Transfer RNA (tRNA) molecules share conserved features that enable proper processing and charging, promoting interactions with mRNA and the ribosome to perform their essential role as the central adaptor molecule of translation. Yet despite these shared functional constraints, cellular tRNA repertoires are remarkably heterogeneous. At the primary sequence level, most organisms encode multiple tRNA isotypes that are charged with the same amino acid to decode distinct mRNA anticodons (isoacceptors), and many species also encode multiple, seemingly redundant isodecoders that share the same anticodon but contain additional sequence differences. The number of isodecoder copies present in the genome and the degree of sequence divergence within isodecoder families vary across evolutionary clades (Chan and Lowe 2015), and not all isodecoders are actively expressed. Furthermore, mitochondria and chloroplasts encode tRNA genes within their genomes, while also importing at least some nuclear-encoded tRNAs from the cytoplasm (Salinas-Giegé et al. 2015). This situation is further complicated in metazoans by tissue-specific differences in tRNA expression (Dittmar et al. 2006), and one recent study found tissue-specific differences in expression and aminoacylation of mammalian mt-tRNAs (He et al. 2021). In addition, many bacterial and eukaryotic viruses also encode tRNA genes within their own genomes despite predominantly relying on the host translational machinery during infection.

RNA modifications confer an additional layer of complexity to the study of tRNA biology. Mature tRNAs contain multiple post-transcriptional modifications per molecule, and the frequency of modification across tRNA nucleotides ranges from 6.5% to 16.5% for bacterial, archeal, eukaryotic, and organellar tRNAs, with nuclear-encoded tRNA from budding yeast containing an average of 13 modifications per tRNA (Phizicky and Hopper 2023). These modifications are abundant and diverse; more than 90 distinct chemical modifications have been identified in tRNA molecules (Cappannini et al. 2023) and demonstrated to affect tRNA folding, structural stability (Whipple et al. 2011; Lorenz et al. 2017), translational fidelity and efficiency (Liu et al. 2022), and the generation of tRNA-derived fragments (Lyons et al. 2018). Moreover, a growing body of evidence highlights the importance of tRNA modifications in human health and disease (Suzuki 2021).

Nanopore sequencing has enabled medium to high throughput interrogation of bacterial and eukaryotic tRNA sequences, as well as insights into select tRNA modifications (Thomas et al. 2021; White et al. 2023; Lucas et al. 2023; Sun et al. 2023). Nanopore sequencing has been commercially developed by Oxford Nanopore Technologies (ONT), which released its first direct RNA sequencing chemistry in 2017 (Garalde et al. 2018). By capturing current changes over time as tRNA molecules are fed through engineered biological nanopores by a specialized helicase, this approach permits the simultaneous identification and quantification of individual tRNA species by sequence, as well as detection of signal disturbances and/or sequence “errors” consistent with modified ribonucleotides (White and Hesselberth 2022). Direct sequencing of native RNA also removes a major technical obstacle to tRNA sequencing, as cDNA synthesis of tRNA is complicated by their extensive modifications and secondary structure (Padhiar et al. 2024).

Detecting RNA modifications in nanopore sequencing relies on the principle that nucleotide modifications disrupt the flow of ions through the nanopore and/or the speed at which modified vs. unmodified RNA passes through the pore (White and Hesselberth 2022); thus, changes to either the helicase or the nanopore have the potential to alter modification signals. While a number of RNA modification types have been detected using nanopore sequencing, a recent update to ONT’s sequencing chemistry motivates re-evaluation of signals at sites of RNA modification. The most recent generation of direct RNA sequencing (RNA004) contains a faster helicase (averaging 130 vs. 70 base pairs / second), and samples are sequenced on specific flow cells containing engineered nanopores that are distinct from their predecessors. These changes have been touted to generate throughput and accuracy improvements for direct RNA sequencing, and were accompanied by a modified basecalling model for m6A detection. However, it remains to be seen how changes to ONT direct RNA chemistry impact detection of other RNA modifications, including those that produce specific signals generated via interactions between the modification and the helicase protein (Stephenson et al. 2022; Burrows and Fleming 2023; White et al. 2023).

In this study, we leveraged the diversity of tRNA repertoires and modifications to evaluate nanopore signals from (i) tRNA from multiple organisms and (ii) at known sites of tRNA modifications characterized previously by orthogonal methods. We sequenced tRNA from six species across diverse evolutionary clades using ONT’s current and previous direct RNA sequencing chemistries, enabling us to compare the reproducibility of tRNA abundance metrics across both methods. We found that while most tRNA modifications produce detectable error signals, the magnitude of these signals varies depending on the selection of both sequencing chemistry and basecalling model. Libraries produced using the RNA004 chemistry generated ∼10-fold higher yields than their first-generation counterparts, enabling us to exploit this additional sequencing depth to detect tRNAs of non-nuclear origin within a larger cellular tRNA repertoire. We demonstrate detection of mitochondrial-specific modification signals in a model eukaryote and find evidence for changes in tRNA repertoires and bacterial tRNA modifications during bacteriophage infection. This comprehensive examination of the impact of distinct RNA modifications on nanopore sequence data sets the stage for researchers to develop more sophisticated machine learning approaches to detect individual tRNA modifications by nanopore sequencing.

## Results

### Updates to direct tRNA library preparation improve sequencing yields

To evaluate the suitability of the most recent ONT direct RNA chemistry and flow cells to nanopore tRNA sequencing, we established a standardized, streamlined protocol for generating tRNA libraries. Based on the strategy developed by (Thomas et al. 2021) and extended in (Lucas et al. 2023), we isolate total RNA from multiple species, treat with base to deacylate tRNA molecules, select for RNAs 17-200 nt in length, and then ligate double stranded RNA adapters specific to the structure, sequence, and end chemistry of tRNA molecules. After a bead purification, these ligation products are next ligated to the ONT-supplied RTA adapter, quantified, and then ligated to helicase-loaded adapters (**Fig. 1A**). Two key refinements in our protocol are the use of commercially available SPRI beads optimized for tRNA purification, as tRNA molecules are not efficiently bound by most SPRI beads (**Fig. S1A-C**), and the optimization of molar ratios for all ligations to reduce carry-through of free adapters into sequenced libraries (**Fig. S1D**). As in (Thomas et al. 2021), we also omit reverse transcription, as (Lucas et al. 2023) only reported a ∼1.4-fold increase in library yields with this step. Together, these changes enable the protocol to be completed in approximately five hours from the first ligation step to loading the library onto flow cells for sequencing.

**Figure 1.**
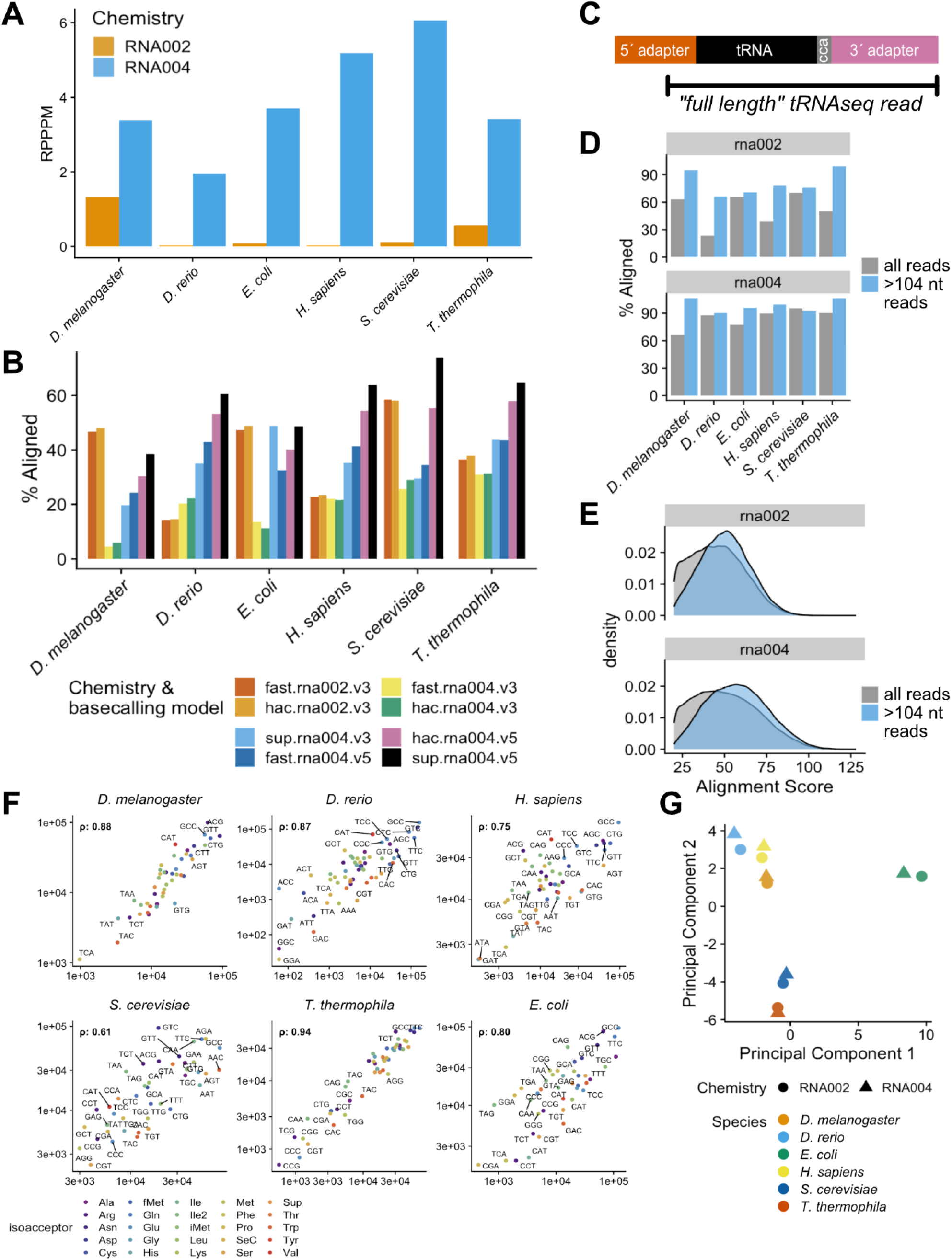
Second generation (RNA004) nanopore direct tRNA sequencing produces higher alignments and sequencing yields with comparable capture of tRNA repertoires. (**A**) Comparative throughput across first (RNA002) and second (RNA004) generation direct RNA chemistries as measured in reads per pore per minute (RPPPM) shows an average 10 fold higher yield for second generation tRNA sequencing libraries. (**B**) RNA004 libraries basecalled with the most recently released and highest accuracy Dorado basecaller (*sup v5*) have the highest alignment rates across all species sequenced with both directly RNA chemistries. (**C**) Schematic outlining the strategy for post-alignment filtering to identify “full length” reads from nanopore tRNA sequencing. (**D**) Lower alignment rates for all reads (gray) vs. reads longer than 104 nucleotides (blue) motivate application of the post-alignment filtering step above. (**E**) Reads ≥105 nt in length exhibit higher alignment scores in both first- and second-generation direct tRNA sequencing libraries. (**F**) Correlation between tRNA isodecoder abundances in RNA002 and RNA004 libraries prepared from *D. melanogaster*, *D. rerio*, *E. coli*, *H. sapiens*, *S. cerevisiae* and *T. thermophila* tRNA. (**G**) Principal components analysis of tRNA repertoires (measured by isodecoder abundance scores), with organism indicated by color and sequencing chemistry by shape, showing that libraries cluster by species and not by sequencing chemistry.

We generated matched pairs of tRNA sequencing libraries from five eukaryotic species and one model prokaryote on the previous (RNA002) and current (RNA004) generation direct RNA sequencing kits from ONT. Each sample was prepared using an input of 300 ng of isolated small RNA as input (containing approximately 10 picomoles of tRNA), and then split in half for RNA002 and RNA004 workup after the RTA ligation step. Equimolar quantities of RTA-ligated material were then ligated to the helicase-loaded RMX (RNA002) or RLA (RNA004) adapters (Fig. 1B), which we estimate are provided at a concentration of 50 nanomolar based on radiolabeling and spectrofluorometric quantation (**Fig. S1E**). RNA002 libraries were loaded onto *R9.4.1* flow cells and sequenced using a custom MinKNOW configuration to prevent the sequencing software from discarding short reads (Lucas et al. 2023), while RNA004 libraries were loaded onto *RNA* flow cells and sequenced using the default direct RNA settings in MinKNOW 23.07, which incorporate the configuration changes described for RNA002 above. Libraries were sequenced on a combination of MinION Mk1b and PromethION 2 Solo instruments, with flow cells serially washed and re-loaded due to the current absence of barcoding solutions for RNA004 multiplexing.

tRNA sequencing libraries prepared using the SQK-RNA004 kit yielded an average of 10-fold increase in sequencing reads compared to the same samples prepared using the first generation sequencing chemistry. To generate a meaningful comparison between sequencing runs performed on different ONT instruments and at different points in the lifetime of a nanopore flow cell, we calculated a normalized yield metric, *reads per pore per minute* (RPPPM) for all libraries. We define RPPPM as the total number of reads generated during the run divided by the length of the run in minutes, which is further divided by the number of pores available at the start of the sequencing run. This metric compensates for the fact that the number of pores available for sequencing directly impacts sequencing yield and the percentage of active pores decreases over the lifetime of a nanopore flow cell. RNA004 libraries displayed a mean RPPPM of 3.49, compared to 0.36 for RNA002 libraries (**Fig. 1A** and **Table S1**). While this metric can be depressed by extended sequencing run times (which we previously observed were often necessary to obtain adequate yields from RNA002 libraries) the approximate 10-fold increase in mean RPPPM observed across these libraries is consistent with our initial pilot experiment, where parallel 30-minute sequencing runs of budding yeast tRNA on newly-opened PromethION flow cells yielded 21,459 RNA002 and 204,731 RNA004 reads, respectively (**Table S1**).

### Read filtering and basecaller model selection impact tRNA alignment

Nanopore basecalling algorithmically converts raw electrical signal data in the form of current intensity over time into nucleotide sequences. We used Dorado to basecall our sequencing data, which is integrated into MinKNOW, the operating software for ONT sequencing instruments, and can perform basecalling in real time during the sequencing run. As ONT’s first and second generation direct RNA sequencing kits use distinct helicases and protein nanopores, samples prepared using these two chemistries must be basecalled using separate algorithms trained on the respective signals from each. To compare our identical input samples prepared using these two chemistries, we re-basecalled all libraries using all available sequencing models in Dorado 0.7.0, and aligned the resulting data using BWA MEM. We found that basecalling with the direct RNA version 5.0 super accuracy (*sup v5*) Dorado model produced the highest alignment rates for all RNA004 libraries (**Fig. 1B** and **Table S2**).

While RNA004 libraries basecalled with the *sup v5* model outperformed other chemistry and basecalling conditions for all samples, we observed some variability in the impact of basecalling model selection and chemistry on the magnitude of improvement in tRNA alignment rates across different species. This could be due to experimental variability or intrinsic, species-specific differences in the population of tRNA molecules (such as their structure, modifications, or processing) that could affect tRNA sequencing and/or basecalling. In particular, *D. melanogaster* and *E. coli* tRNA libraries showed only modest improvements in alignment rates between RNA002 and RNA004 chemistry.

Nanopore tRNA sequencing was first performed by (Thomas et al. 2021) on the RNA002 chemistry in *E. coli*, with additional refinement of sequencing parameters and application to eukaryotic species by (Lucas et al. 2023) and our own group (White et al. 2023). All of these studies used BWA MEM to perform heuristic local alignments of tRNA reads, using a set of relaxed parameters for alignment to improve the mappability of short, error-prone nanopore reads. While (Lucas et al. 2023) empirically tested these parameters to minimize false positive alignment to tRNA antisense sequences, evaluation of the accuracy of tRNA alignments from nanopore sequencing has been limited, and is complicated by the evolving accuracy of direct RNA sequencing itself. Recent work using first generation nanopore tRNA sequencing to examine queuosine-modified tRNAs raised questions about false positive alignments when performing heuristic local alignments as in the studies above, and suggested BWA MEM may produce off target or low quality alignments of nanopore tRNA sequencing (Sun et al. 2023). This motivated us to reevaluate our approach to tRNA alignment in this study.

In these and previous nanopore tRNA sequencing using RNA002 chemistry, we observed a substantial proportion of reads with lengths shorter than expected for mature, adapter-ligated tRNA molecules (**Fig. S2A,B**). While we can only speculate on whether these reflect genuine smaller RNA species captured in these libraries or reads artificially truncated during basecalling, regardless of their origin, shorter sequences have an intrinsically higher probability of aligning to the incorrect tRNA isodecoder. We therefore added a post-alignment length filtering step to all libraries, requiring alignments to span the full length of the tRNA body, plus the entirety of the 3′ tRNA adapter, and 9 nucleotides into the 5′ adapter (to reflect the expected sequence basecalled before the helicase loses contact with the 5′ end of the nucleic acid during sequencing) (**Fig. 1C**). As mature tRNA sequences vary in length, this equates to alignments spanning 108 - 143 nucleotides of reference sequence for the organisms sequenced in this study.

Filtering for mature, full length tRNA reads boosts both alignment rates and alignment scores using BWA MEM. When we apply this filtering step to tRNA sequencing libraries generated using the highest accuracy basecalling model for each chemistry, we retain a median 43% of reads in RNA002 libraries and 62% of reads from RNA004 libraries. To explore the impact of this post-alignment filtering step, we chose a length cutoff of 105 nt and examined the percent of reads aligning above and below this length threshold. While only 56.8% of unfiltered reads in RNA002 libraries align to a tRNA reference, reads ≥105 nt in length have a 77.2% median alignment rate. RNA004 libraries display a similar, if less dramatic increase in alignment rates by length, going from a median 88.6% alignment rate for all reads to a median 97.9% of ≥105 nt reads aligning (**Fig. 1D**). After post-alignment full length read filtering, the distribution of alignment scores shifts higher with both chemistries (**Fig. 1E**). We further evaluated the impact of this post-alignment filtering strategy on reads basecalled with all available Dorado models (**Fig. S2C,D**), and concluded that the combination of full length filtering and highest accuracy models (*hac v3* for RNA002 or *sup v5* for RNA004) produce the best combination of high scoring alignments while retaining sufficient reads for downstream analysis. Unless otherwise indicated, all data in the remainder of this manuscript has been pre-processed accordingly.

### tRNA abundance correlates across first and second generation direct RNA sequencing chemistries

In light of the limited changes to library preparation between SQK-RNA002 and SQK-RNA004, we hypothesized that second generation direct RNA sequencing would produce comparable capture of individual tRNA species to the previous approach. We compared normalized tRNA abundances for all libraries, and observed that Pearson’s correlation coefficients between RNA002 and RNA004 libraries ranged from 0.61 to 0.94 (**Fig. 1F**). Thus, despite changes to the helicase, nanopore, and basecalling algorithms, tRNA abundance measurements obtained from both nanopore direct RNA sequencing approaches (RNA002 and RNA004) exhibit a stronger correlation with each other than was previously reported for RNA002 vs. Illumina-based tRNA sequencing methods (Lucas et al. 2023). Moreover, a principal components analysis of normalized tRNA counts from all libraries reveals that these samples cluster more closely by species than by library chemistry, underscoring the robustness of this approach for capturing biologically relevant variations in tRNA abundance (**Fig. 1G**).

### >90% of known tRNA modifications are detectable by basecalling error in RNA004 sequencing

tRNA are densely packed with chemical modifications that impart both structural and functional attributes, with implications for tRNA folding, stability, turnover, charging and decoding. RNA modifications are a known source of nanopore signal distortion, and can produce basecalling “error” signals in the form of mismatches, in which the basecaller assigns the wrong nucleotide, insertions, in which the basecaller calls an additional nucleotide, or deletions caused by the failure to call any base at all (Liu et al. 2019; Parker et al. 2020; Price et al. 2020; Jenjaroenpun et al. 2021; Begik et al. 2021). Although many groups leveraged first generation nanopore direct RNA sequencing to identify select RNA modifications (White and Hesselberth 2022), the field has not yet settled on a standard approach for RNA modification detection from direct RNA sequencing data. While the technology can in principle identify many types of modifications within the same molecule, in practice, a number of technical challenges need to be addressed.

Previous studies using RNA002 chemistry used a combination of computational approaches to analyze modification signals. In general, these approaches involved the identification of candidate modified sites by changes in basecalling error at the sequence level, followed by an optional analysis of raw signals (current x time data or “squiggle”) at known or candidate modification sites. As modification-dependent signals are produced via specific interactions as the modified nucleotide is transiting through the helicase/pore architecture, the shift to RNA004 chemistry requires a re-analysis of all RNA modifications previously analyzed by nanopore to determine whether modification signatures characterized using the RNA002 chemistry remain consistent or change with the introduction of a different helicase and nanopore protein.

tRNAs offer a unique opportunity to examine the signals produced at a large number of previously validated, high stoichiometry RNA modification sites across a range of organisms, RNA molecules, and sequence contexts. The MODOMICS database catalogs 189 distinct modifications identified across coding and non-coding RNAs, of which 67 have been observed on tRNAs. Across the five eukaryotes, one prokaryote, and one bacteriophage genome sequenced in this work, 55 unique modifications have been observed on at least one tRNA isodecoder (Cappannini et al. 2023). To evaluate nanopore signals produced across a range of tRNA modifications, we analyzed basecalling error signatures on all sequenced tRNAs with annotated sites of RNA modifications in MODOMICS. We filtered for sites of known modifications with at least 30 mapped tRNA reads in our RNA004 sequencing data, yielding 43 types of tRNA modifications observed at >3700 unique locations, spanning 396 distinct 5mer sequence contexts.

Ribonucleotides at known positions of tRNA modifications produce larger basecalling error signals than positions that lack modification information, with the largest magnitude difference seen at modified adenosines, which have a 51% higher basecalling error frequency than adenosines lacking annotated modifications in MODOMICS. Uracil and guanosine modifications also display >25% increases in basecalling error frequency, as opposed to cytosine modifications, which have a median increase of 3% compared to their unmodified counterparts (**Fig. 2A**). Across the board, mismatches were the primary driver of these basecalling error differences, followed by deletions and then insertion frequency (**Fig. S3A-D**).

**Figure 2.**
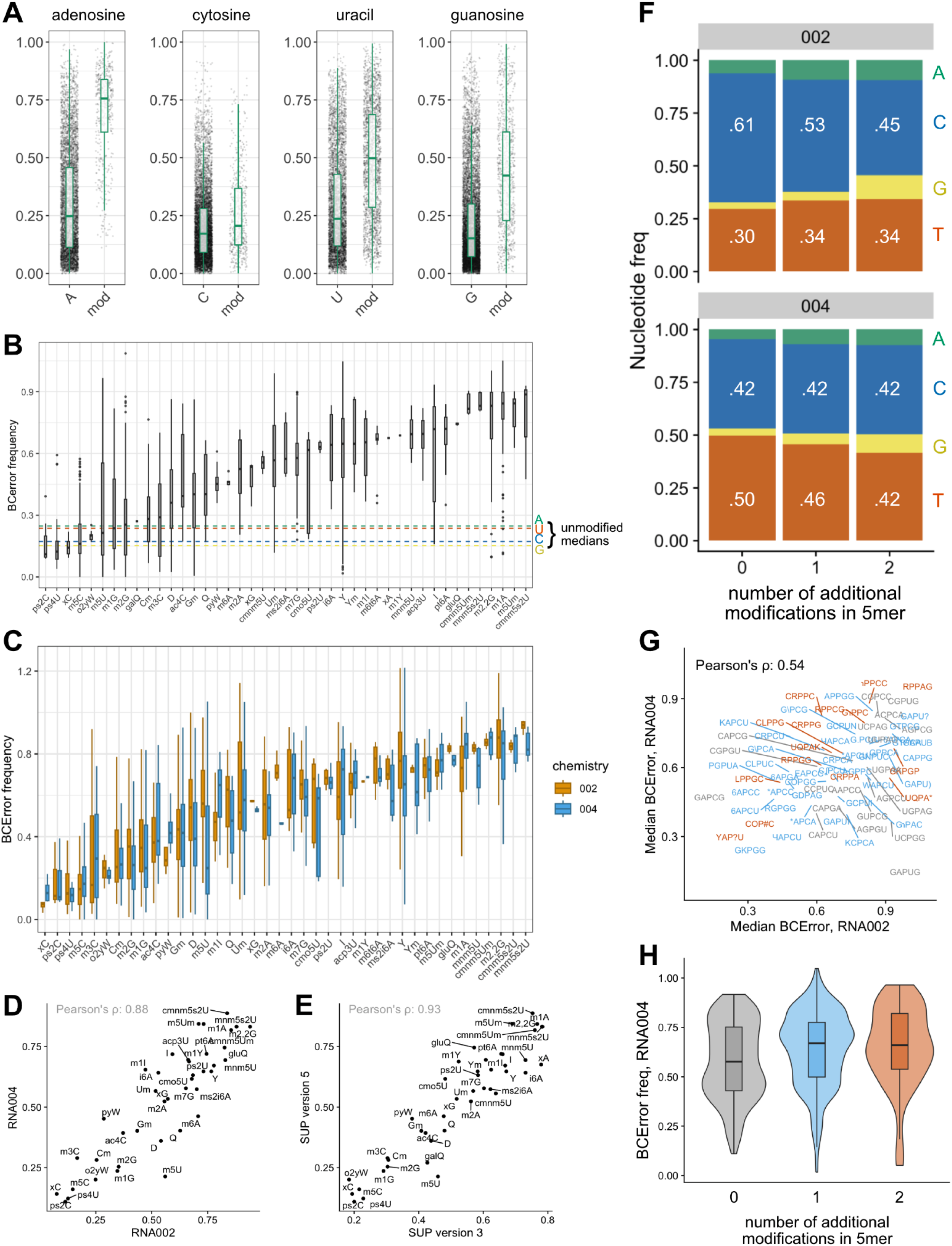
Over 90% of known tRNA modifications with ≥30X coverage in RNA004 sequence data generate basecalling signals above background error threshold; however, the magnitude and consistency of these signals depends on the modification in question, the sequencing chemistry, and the basecalling model selected. Modification abbreviations used in this figure are derived from MODOMICS and can be found in Table S3. (**A**) Basecalling error frequencies (summed insertion, deletion and mismatch frequencies) at known tRNA modifications annotated in the MODOMICS database (mod) vs. unmodified A, C, U and G nucleotides in RNA004 sequencing data. Each dot represents a position on a tRNA isodecoder with at least 30 reads aligning. (**B**) 39 of 43 RNA modifications with ≥30X coverage in our RNA004 sequencing libraries produce basecalling error frequencies above the background threshold for the corresponding unmodified nucleotide. Modifications are ordered from left to right from lowest to highest median error frequency, with the background levels for unmodified nucleotides in panel **A** indicated via dashed lines. (**C**) Signals produced by some RNA modifications vary dramatically depending on sequencing chemistry. RNA002 data is indicated in orange, and second-generation RNA004 data is plotted in blue. Note that for the previous chemistry, select positions have basecalling error frequencies exceeding 1 due to high rates of insertions and deletions. (**D**) The median basecalling error frequencies for all tRNA modifications in panel **C** have a 0.88 Pearson’s correlation coefficient between RNA002 and RNA004 libraries. (**E**) When RNA004 data is basecalled using two different versions of the Dorado super high accuracy (‘sup’) model, the tRNA modifications plotted in panel **B** have a higher Pearson’s correlation coefficient (0.93) than libraries produced using different chemistries. (**F**) Known sites of tRNA pseudouridylation display variable levels of U-to-C mismatching in first- and second-generation direct RNA sequencing libraries, with additional differences depending on the number of neighboring modifications present within a 5 nucleotide window surrounding the pseudouridine. (**G**) The median basecalling error at pseudouridylated positions is highly variable depending on both sequencing chemistry and surrounding sequence context. Individual 5mer sequence contexts are plotted in different colors depending upon the number of additional modifications present within the 5mer, using the same color scheme as in the panel below. (**H**) In RNA004 libraries, known pseudouridylated sites display higher basecalling error frequencies on average as additional modifications are present within the same 5 nucleotide window.

We further analyzed our data at the level of individual modifications, allowing us to compare the relative magnitude of basecalling error signals observed at abundant tRNA modifications. **Figure 2B** shows all 43 tRNA modifications present in our RNA004 data after *sup v5* basecalling and filtering, ordered from lowest to highest median basecalling error frequency on the X axis, with the median basecalling error scores for unmodified nucleotides from panel **A** indicated in dashed horizontal lines. (For a complete set of abbreviations for all modifications evaluated, see **Table S3**.) All but four modification types analyzed exceeded the median basecalling error score for their corresponding unmodified nucleotide. In these plots, each dot represents a single tRNA isodecoder’s sequence context with at least 30 reads aligning. We observe an overall trend towards bulkier modifications producing higher basecalling error values, with some types of modifications producing a wide spread of basecalling error frequencies across different tRNA isodecoders and sequence contexts (e.g., dihydrouridine [D], pseudouridine [Y], and inosine [I]), while other abundant RNA modifications (for example, m1A) yield a much tighter distribution of values.

### Most tRNA modification signals closely correlate across first and second generation direct RNA sequencing chemistries

Figure 2C compares the distribution of basecalling error scores between the RNA004 data described above and the matched first-generation chemistry libraries basecalled with Dorado’s RNA002 *hac* model. While the median basecalling error for all modifications are closely correlated across the two chemistries (Fig. 2D), the magnitude and spread of these values vary depending on the modification examined. Ten tRNA modifications (including 5-methyluridine, 6-methyladenosine, queuosine, dihydrouridine and pseudouridine) exhibited a >10% reduction in median basecalling error score between RNA002 and RNA004 libraries, with while seven more (including inosine, 1-methylinosine, and 1-methyladenosine) instead displayed a more than 10% increase with the second-generation chemistry. Finally, three additional types of tRNA modifications only achieved ≥30X coverage in our RNA004 libraries, consistent with the higher throughput of the second generation direct RNA chemistry. In addition to these throughput differences, it is worth noting that lower basecalling accuracy and alignment rates in our first generation tRNA sequencing libraries may limit the ability to make accurate comparisons of tRNA modification signals across these two approaches (see **Fig. S3E** for background basecalling error rates in RNA002 tRNA sequencing).

Further comparison of basecalling error at known tRNA modification sites under different basecalling conditions revealed close correlation of sequence-level error signals across RNA004 basecalling models. Fig. 2E displays median basecalling error signals at various RNA modifications for the same libraries basecalled with two different versions of Dorado’s *sup* model: the original version 3 model released alongside the RNA004 chemistry and the recently released version 5 update, which uses a distinct, transformer-based model architecture. Both these comparisons and version 5 *sup* vs. *hac* basecalling (data not shown) are more tightly correlated, indicating that tRNA modification signals are most impacted by the change in library chemistry, but that the choice of basecalling model selection may also have implications for basecalling error signal depending upon the modification in question.

### Basecalling error signals at pseudouridylated positions vary across sequencing chemistry

U-to-C miscalling is a well characterized hallmark of pseudouridylation in nanopore sequencing data for libraries generated using the first generation RNA002 chemistry (Begik et al. 2021). We observed a 12% drop in median basecalling error from RNA002 to RNA004 libraries at annotated pseudouridine sites (Fig. 2C), which led us to explore how conserved this signature is in the new nanopore direct RNA chemistry, as well as the possible contributions of neighboring RNA modifications on pseudouridine signals. Figure 2F compares the rate of miscalling (one component of basecalling error) between RNA002 and RNA004 libraries over 5mer sequence contexts with a central pseudouridine where either 0, 1, or 2 additional RNA modifications are annotated in MODOMICS within the same region. When we examine mismatching alone, sites with no additional modifications were miscalled as cytosine 61% of the time in RNA002 libraries, whereas in RNA004 libraries, this rate drops to just 42%. While both chemistries show a stepwise increase in U-to-A and U-to-G mismatches when additional modifications are present within the 5mer, for RNA004 libraries, the rate of U-to-C mismatches remains a constant 42% independent of neighboring modifications, while the presence of an additional modification reduces the frequency at which the basecaller correctly calls pseudouridylated sites as uridine by 4% per additional modification within the 5mer. By contrast, RNA002 libraries show a stepwise reduction in U-to-C mismatching as neighboring modifications are added.

Although the presence of neighboring RNA modifications clearly influences the likelihood that a pseudouridylated site will be miscalled, the rate of basecalling error across such sites remains highly heterogeneous and only weakly correlated between RNA004 and RNA002 libraries. Individual sequence contexts have a Pearson’s correlation coefficient of 0.54 between the two chemistries, with no obvious trend based on the number of additional modifications present within the 5mer (Fig. 2G). Indeed, the basecalling error frequencies for pseudouridylated sites with 0, 1, or 2 additional modifications in RNA004 libraries have large overlaps in their distributions, suggesting that sequence context remains a major contributor to the magnitude of error signal observed at this modification (Fig. 2H). Finally, we also note that 5-methyluridine (5mU) modifications, which are commonly found one nucleotide upstream of pseudouridine at position 55, display even larger differences between RNA002 and RNA004 libraries (a 35% drop in basecalling error signal, see **Fig. 2C and S3F**). As different types of RNA modifications can produce different “spreads” of basecalling error (Begik et al. 2021; White et al. 2023), disentangling the signals from adjacent modifications, especially in the context of modification interdependencies like those described for tRNA anticodon and T loops (Han and Phizicky 2018; Barraud et al. 2019), remains challenging even as direct RNA sequencing accuracy improves.

### Direct RNA sequencing enables detection of mitochondrial tRNA

High throughput studies of mitochondrial tRNA (mt-tRNA) biology have been limited by the low abundance of mt-tRNA species relative to nuclear-encoded tRNAs. We sought to evaluate the suitability of nanopore direct tRNA sequencing to study mitochondrial tRNAs, using the budding yeast *Saccharomyces cerevisiae* as a model. *S. cerevisiae* encodes 24 tRNA genes in its mitochondrial genome, providing complete decoding capabilities within the mt-tRNA repertoire via wobble decoding. Despite this, *S. cerevisiae* mitochondria also import select nuclear-encoded lysine and glutamine tRNAs from the cytoplasm (Martin et al. 1979; Rinehart et al. 2005). Under fermentable conditions, respiration is dispensable in *S. cerevisiae*, allowing cells to survive in the absence of mitochondrial function (Ephrussi 1953). Such mutants, termed “petite” due to their slow growth and small colony phenotype, can be induced at higher frequencies via treatment with ethidium bromide (EtBr) or other drugs, leading to extensive deletion (⍴^-^) and even complete loss (⍴^0^) of mtDNA (Slonimski et al. 1968; Contamine and Picard 2000).

We generated petite colonies of *S. cerevisiae* with ethidium bromide, isolated small RNA, and generated tRNA sequencing libraries using the RNA004 chemistry, enabling us to compare the number of reads aligning to mitochondrial tRNAs between petite yeast vs. wild type (“grande”) yeast. After alignment to a tRNA reference containing all tRNA isodecoders from both the nuclear and mitochondrial (Turk et al. 2013) yeast genomes, we observed an 85% reduction in mitochondrial-derived reads in petite compared to grande yeast, while the amount of nuclear tRNA recovered showed a 0.4% global increase (Fig. 3A, B). While the remaining mitochondrial tRNA reads in our petite strain could be consistent with residual mitochondrial DNA after EtBr treatment, another possibility is that sequence similarities between mitochondrial and nuclear encoded yeast tRNAs (**Fig. S4A**) enables some amount of false positive alignment between nuclear tRNA reads and mitochondrial tRNA references.

**Figure 3.**
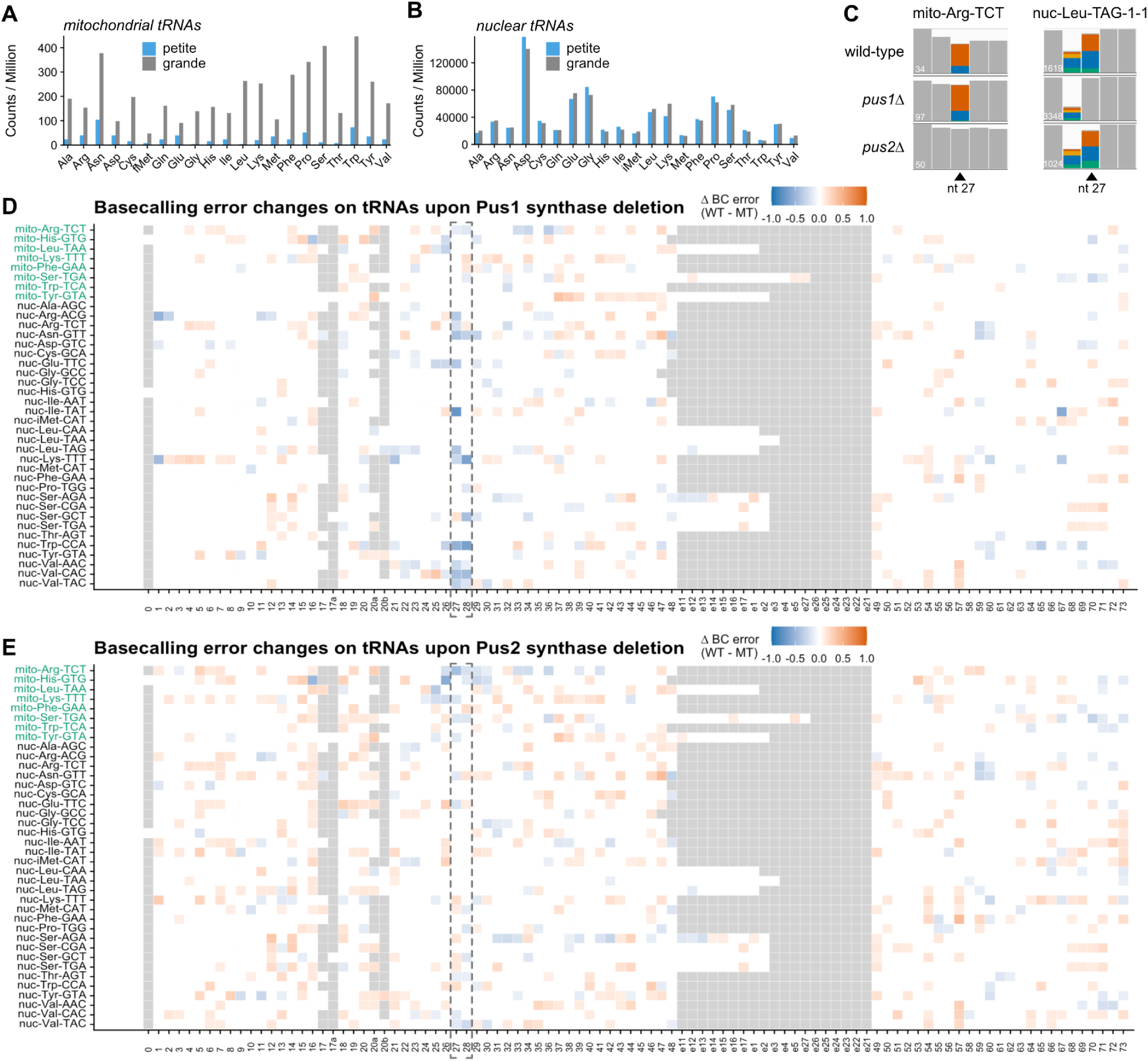
Second-generation nanopore direct RNA sequencing enables analysis of mitochondrial tRNAs and their modifications. (**A**) Counts per million reads aligning to mitochondrially-encoded tRNA isoacceptors from RNA004 tRNA sequencing libraries prepared from budding yeast that were confirmed to be respiration-deficient (*petite*) cells in blue, or their or wild-type (*grande)* counterparts in gray. Petite cells, which have lost their mtDNA upon EtBr treatment, display an 85% reduction in reads aligning to mitochondrial tRNAs. (**B**) Petite and grande cells show commensurate levels of nuclear-encoded tRNAs in RNA004 sequencing data. (**C**) IGV screenshots demonstrate the specificity of tRNA modification induced basecalling error signals to nuclear- and mitochondrial-encoded tRNAs. Deletion of the mitochondrial pseudouridine synthase *PUS2* causes loss of U-to-C mismatching at nucleotide 27 on a representative mitochondrial tRNA (Arg-TCT), but not the nuclear-encoded tRNA Leu-TAG isodecoder. In contrast, deletion of the pseudouridine synthase *PUS1*, which localizes to the cytoplasm, results in loss of U-to-C mismatching at nucleotide 27 on the nuclear-encoded tRNA at the same position. Positions with less than 20% mismatching to the reference base are colored in gray, while colored bars exceed this threshold; the total number of reads aligned to these regions for each library are indicated in white in the bottom left. For Leu-TAG-1-1, the upstream guanosine at position 26 is a known site of N2-dimethylation (**D**) A heat map displaying the change in basecalling error (Δ BC error) between wild-type and *Pus1Δ* cells. Nucleotides 27 and 28, which are pseudouridylated by *PUS1* in the cytoplasm, are highlighted in the dashed box spanning all tRNAs. Mitochondrial tRNAs are indicated in green text at the top of the heat map, while nuclear-encoded tRNAs are in black. A negative Δ BC error value indicates a putative loss of modification; across this biological comparison, these signals are strongest at nucleotides 27 and 28 on nuclear-encoded tRNAs. (**E**) Similar to the heat map in panel **D**, the change in basecalling error between tRNAs isolated from wild-type *S. cerevisiae* and a *PUS2* deletion strain. In the absence of this mitochondrially-localized pseudouridine synthase, we see a drop in basecalling error signal (consistent with loss of pseudouridylation) at nucleotides 27 & 28 that is predominantly restricted to mitochondrial tRNAs. In panels **D** and **E**, not all tRNAs present in the reference are plotted, but rather a representative set of tRNA isodecoders with structural and modification information from MODOMICS are aligned to a shared coordinate set corresponding to the consensus Rfam tRNA structure RF00005. Positions lacking ≥30X coverage or where no nucleotide is present for that particular tRNA isodecoder at the position on the consensus structure are colored gray.

To further confirm that these mt-tRNA aligning reads were genuine, we sequenced additional tRNA libraries from strains with genomic disruptions in the pseudouridine synthase genes *PUS1*, *PUS2*, and *PUS4*. While Pus4 has broad specificity for nucleotide 55 of tRNAs in both the mitochondria and cytoplasm (Becker et al. 1997), Pus1 and Pus2 act at either nucleotide 27 or 28 on a subset of tRNAs, and their targets are further restricted by their subcellular localization. While Pus1 is localized to the cytoplasm, Pus2 is specifically targeted to the mitochondria, where it pseudouridylates a distinct set of tRNA substrates (Behm-Ansmant et al. 2007). As such, we would expect to see loss of pseudouridylation signal at this location on genuine mitochondrial tRNA reads in *pus2Δ* cells, and an analogous loss of signal on nuclear-encoded tRNAs in *pus1Δ* cells.

Our tRNA sequencing data from these three strains is consistent with the localization and activity of each pseudouridine synthase. Figure 3C displays loss of mismatching signal at nucleotide 27 on mitochondrial tRNA Arg-TCT in *pus2Δ* cells, with no change at the corresponding position on a representative nuclear-encoded tRNA. In contrast, deletion of *PUS1* results in a loss of signal at nucleotide 27 on a nuclear encoded tRNA leucine isodecoder (Fig. 3C), while mismatch signals at the upstream guanosine, a known site of N2-dimethylation, remain unchanged. These error signatures are generally consistent when we further plot the change in basecalling error (that is, the combined frequency of insertions, deletions, and mismatches at each position) for additional tRNAs. When *PUS1* is deleted, the largest differences in basecalling error between mutant and wild type are concentrated at positions 27 and 28 in nuclear tRNAs (Fig. 3D), while deletion of *PUS2* produces similar signals on a handful of mitochondrial tRNAs (Fig. 3E). In contrast, when the pseudouridine synthase *PUS4* is deleted, signals at position 27 are unaffected, but mismatches produced by *PUS4*-dependent pseudouridylation at nucleotide 55 are lost on both mitochondrial and nuclear tRNAs (**Fig. S4B**).

### Capture and sequencing of viral tRNAs in a bacteriophage infection model

The study of less abundant tRNA species within a broader biological mixture is also important in the context of viral infection. Despite their reliance on host cell translational machinery for protein synthesis, many bacteriophage (phage) encode tRNAs within their own genomes. Several models have been proposed for how such tRNAs may enable pathogens to evade host defenses and/or bias translation programs during infection (van den Berg et al. 2023; Yang et al. 2021; Bailly-Bechet et al. 2007). While it is clear that phage-encoded tRNAs are targets of host endonucleases and that some phage tRNA modifications and/or phage tRNA sequences may protect phage tRNAs from host endonucleolytic cleavage (Georjon and Bernheim 2023; Ogawa 2016; Ogawa et al. 2021), broader questions about the expression, modification, and aminoacylation of phage tRNAs during infection remain unanswered.

To determine the suitability of second generation direct RNA sequencing to the investigation of these questions, we sequenced tRNA from both untreated *E. coli* or *E. coli* infected with T4 bacteriophage. As before, we filtered for full length reads, and then aligned these sequencing data to a combined reference containing the entire repertoire of *E. coli* tRNA genes, as well as the 7 tRNA genes present in the T4 phage genome. In an uninfected sample, 0.2% of full length reads align to tRNAs from the T4 genome, suggesting a low level of false positive alignment, possibly from sequence similarity between host- and phage-derived tRNAs (compared in **Fig. S5**). After 60 minutes of T4 infection, the number of reads aligning to phage tRNAs increased by 23-fold, compared to a 5% global decrease in host derived tRNAs (Fig. 4A), providing proof of concept for the application of nanopore tRNA sequencing to the study of tRNA in an infection context. Moreover, changes in basecalling error signal on host tRNAs upon T4 infection, including a site-specific decrease in mismatching at nucleotide 55, which is routinely pseudouridylated by the TruB/Pus4 family of pseudouridine synthases to initiate a cascade of downstream modifications within the T loop, raise the possibility that the RNA modification landscape of host tRNAs may be dynamically remodeled in response to phage infection (Fig. 4B). Taken together, these data provide proof of concept for using nanopore tRNA sequencing to study tRNA dynamics in the context of viral infection.

**Figure 4.**
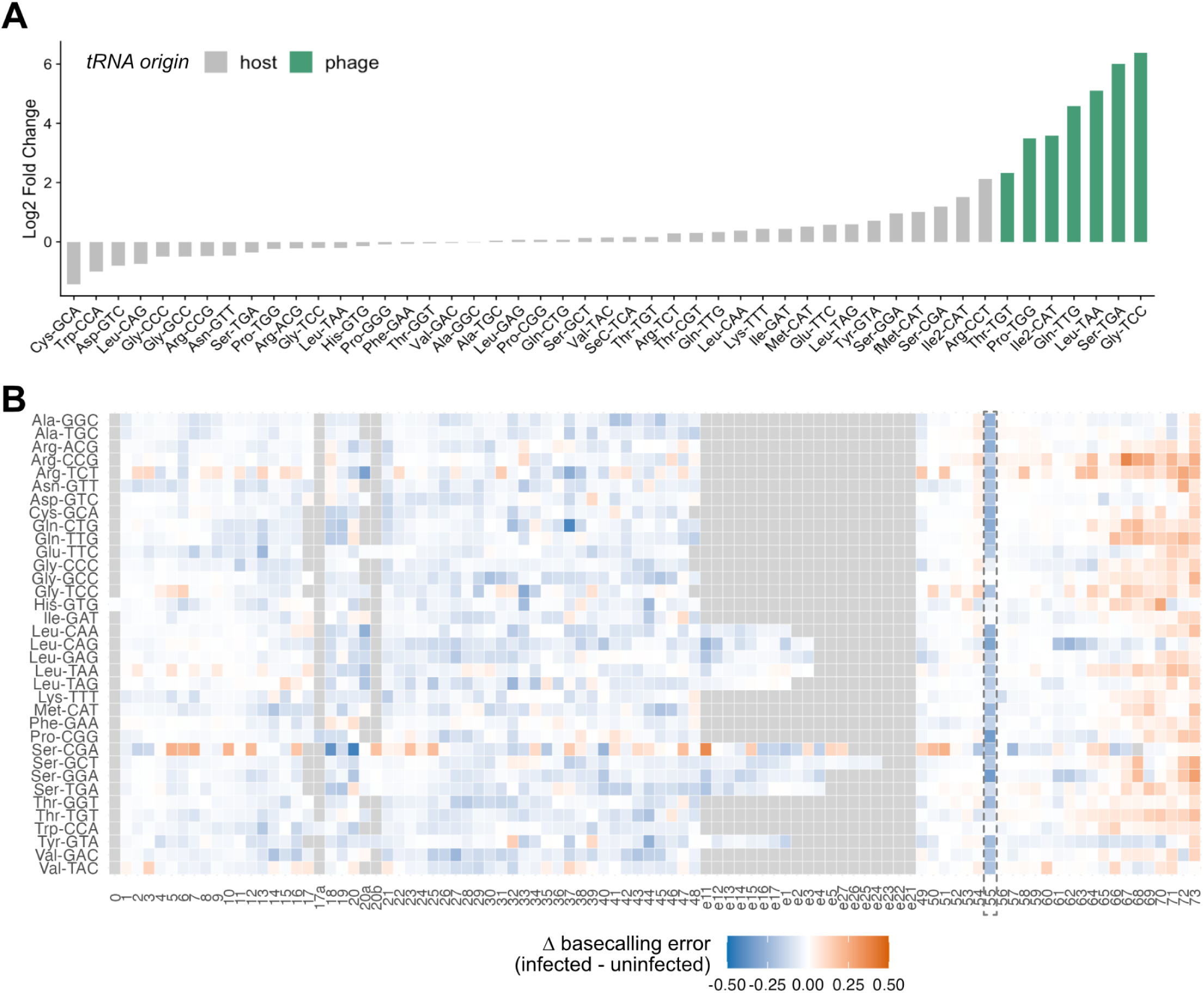
tRNA sequencing of bacteriophage-infected *E. coli* reveals dynamic changes in tRNA abundance and modifications. (**A**) A plot of the log2 fold change in tRNA abundance between uninfected *E. coli* and the same strain after 60 minutes of T4 bacteriophage infection. All reads were aligned to a combined *E. coli* and T4 phage tRNA reference, and phage-encoded tRNAs, which increase the most over this timeframe, are indicated in green. (**B**) A heat map displaying the change in basecalling error on host (*E. coli*) tRNAs between T4-infected and uninfected cells. Negative Δ basecalling error values are indicated in blue, and indicate a putative loss of RNA modification signal upon phage infection, including a dramatic drop in error signal at the known pseudouridylated position 55 (indicated in dashed box). As in Figure 3, positions that lack ≥30X coverage or where no nucleotide is present for that particular tRNA isodecoder at the position on the consensus structure (most notably, in the variable loop) are colored gray on this heatmap.

## Discussion

Nanopore direct RNA sequencing is a promising approach for the study of RNA molecules and their modifications, with major implications for the study of tRNA, the most abundant and diversely modified class of RNAs in cells. However, despite the rapid pace of sequencing technology development, much work remains to enable the identification and analysis of the complete repertoire of tRNA molecules and their modifications. In this study, we evaluate a major update to the Oxford Nanopore direct RNA chemistry via tRNA sequencing across five eukaryotic species and one prokaryote, and demonstrate that these changes, in combination with our own protocol optimizations and refined alignment strategy, enable rapid sequencing of tRNA molecules, greater alignment accuracy, and an order of magnitude higher library throughput than previous approaches. Together, these updates make nanopore tRNA sequencing increasingly tractable for researchers beyond those actively engaged in sequencing method development, with improvements in yield and basecalling accuracy opening up several new possibilities for the study of tRNA biology.

Higher tRNA sequencing yields will enable the study of lowly abundant tRNA species, including organellar tRNAs and tRNAs expressed during infection by pathogens. While our results show only a minor impact of T4 infection on *E. coli* tRNA abundance, future studies will require not only orthogonal validation of putative changes in pseudouridine and/or m5U modifications on host tRNAs, but also examination of how both host and phage tRNA levels throughout an infection time course. Previous work on tRNA dynamics during vibriophage infection indicate that phage tRNA can compensate for degradation of the host translational machinery, including tRNA molecules, though expression of host tRNA may decline and then rebound cyclically if some hosts survive the first round of replication uninfected (Yang et al. 2021). Furthermore, a better understanding of how the composition of the intraorganellar tRNA pool impacts translation may prove invaluable to the study of mitochondrial disease, as mitochondrial dysfunction is implicated in a wide range of disorders caused by mutations in both nuclear and mitochondrial genomes (Russell et al. 2020). Many of these result in myopathies or neurological disorders, and may be caused by mutations in tRNA genes themselves, in aminoacyl tRNA synthetases (aaRSs), in tRNA processing factors, or in RNA modification enzymes. In several of these disorders, an inadequate supply and/or hypomodification of one or more tRNA species has been directly implicated in disease; however, the molecular mechanisms responsible for the resulting tissue-specific phenotypes remain poorly understood (Burgess and Storkebaum 2023), underscoring the need for more comprehensive and high resolution investigations of mt-tRNA pools.

We also expect that higher library throughput and analysis of single reads from nanopore sequencing will facilitate the study of sub-populations of differentially modified tRNA isodecoders. While this study is focused almost entirely on basecalling error signals produced at previously-annotated sites of tRNA modification, ongoing improvements to ONT basecalling models for native ribonucleotides, pseudouridine, and m6A as well as the introduction of new models for additional RNA modifications are expected to facilitate *de novo* detection for an increasing number of epitranscriptomic marks. Indeed, recent updates to ONT basecalling models in Dorado 0.7 now support detection of m6A and pseudouridine modifications in all sequence contexts; however, these models do not include tRNA or similarly complex modified RNA samples in their training data (Oxford Nanopore Technologies, personal communication). While these models were released while this manuscript was in preparation, our initial exploratory analysis using Dorado 0.7 basecalling, reference-anchored pseudouridine detection via Remora, and pileup of high-confidence modification sites using ModKit suggests that v5 basecalling models are only capable of identifying all but highest stoichiometry modified sites on tRNA (e.g., pseudouridylation at nt 55), with high levels of false positives and current limited utility to per-read analysis (data not shown). Given the rapid evolution of tools and basecalling models, researchers should take care to validate new approaches for modification identification, particularly for tRNA modifications where model training data may not sufficiently capture the surrounding sequence, modification, and structural context, all of which may influence raw nanopore signals, and by extension, modification calling.

Our analysis of many tRNA isodecoders from a range of organisms has identified a collection of tRNA modifications that produce dramatic distortions in raw signals from nanopore sequencing capable of confusing the Dorado basecaller, generating basecalling errors. While not every modification type produces the same type or magnitude of basecalling error in all sequence contexts, the modifications with a robust error signature identified in Figure 2B represent low hanging fruit for future training efforts towards *de novo* identification of additional RNA modifications from RNA004 sequencing data, as well as modifications with less robust and/or more heterogeneous whose identification may require more intensive model training. *De novo* identification of the complete set of RNA modifications via nanopore sequencing and other approaches is an area of great interest, but the need to develop ground truth standards for modifications in different stoichiometries and sequence contexts makes this a grand challenge for the field (Begik et al. 2022; National Academies of Sciences et al. 2024). However, since tRNAs’ core biological function imparts constraints upon their structure, sequence, and modification status, we note that generating a complete repertoire of sequence contexts may not be necessary when training models specifically to interpret tRNA modification signals. Our future efforts will leverage these tRNA sequencing data, along with genetic and *in vitro* transcribed controls, to develop strategies to identify subpopulations of differentially modified tRNAs from raw nanopore signal, as well as to investigate coordination of tRNA modifications during their complex maturation process.

## Materials and methods

### Sample-specific genotypes and growth conditions

#### Bacterial growth and phage infections

*Escherichia coli* strain MG1655 was grown in 5 mL LB media at 37°C overnight. The next day, 50 mL LB media was inoculated with the overnight culture to an OD_600_ = 0.025 and allowed to grow to an OD_600_ = 0.3. Once desired OD_600_ was reached, cultures were infected with T4 bacteriophage at an MOI of 0.25 and allowed to adsorb for 10 minutes. Infected cultures were placed at 37°C for 60 minutes. Total RNA isolation was by acid phenol:chloroform extraction.

#### Yeast

The *Saccharomyces cerevisiae* parental strain (BY4741), as well as *pus1Δ*, *pus2Δ* and *pus4Δ* knockouts were obtained from Yeast Deletion Collection (Open Biosystems). Petite cells from were generated from an S288C background (MATa/alpha ho/ho ura-352/ura-d) by adding ethidium bromide to yeast extract, peptone (YEP) media with 2% glucose at a final concentration of 25 µg/mL and incubating with rotation at 30°C overnight. The culture was diluted 1:10,000 and plated on YEP glucose solid media for single colonies. Colonies were allowed to grow for 72 hours before inspection to identify petite phenotypes. Candidate colonies were streaked onto YEP glucose media and patched onto yeast extract, peptone, glycerol, ethanol (YPGE) solid media to confirm their inability to respire.

To isolate RNA, single colonies were inoculated into 10 ml YEP glucose cultures and incubated at 30°C overnight with rotation. The next day, cultures were centrifuged and total RNA isolated by hot phenol extraction.

#### Tetrahymena

*T. thermophila* strain B1868 was grown in SSP media (2% peptone, 0.1% yeast extract, 0.2% glucose, 33 µM FeCl3), shaking at 30° C until cell density reached 10^5^ cells / mL. Cells pellets were harvested by centrifugation and frozen at -80° C prior to total RNA extraction with acid phenol:chloroform.

#### Human cell culture

Hs27 human fibroblast cell cultures (CRL-1634) were handled according to the instructions of the manufacturer. In brief, cells were plated at a density of 7,500 cells per cm2 using DMEM media supplemented with 10% fetal bovine serum (FBS) and 1% antibiotic. Cell cultures were passaged upon 80% confluency and only passages between 3-8 were used for subsequent RNA extractions. For harvesting, cell cultures were washed with 1X phosphate buffered saline - pH 7.4 (PBS, 10010031, Gibco), trypsinized using Trypsin-EDTA (25200056, Gibco) for five minutes at 37°C, and then collected by centrifugation at 700 rcf. Pelleted cells were further washed using PBS and total RNA was extracted using TRIzol (15596026, Invitrogen) and isopropanol precipitation.

#### Insect cell culture

Schneider 2 Drosophila melanogaster cells (R69007, ThermoFisher Scientific) were plated and incubated according to the instructions of the manufacturer. In brief, cell cultures were maintained in Express Five™ SFM medium (10486025, ThermoFisher Scientific) supplemented with 10% L-glutamine. Cell cultures were passaged or harvested for RNA extraction once they were 90-100 % confluent. Cells were collected by centrifugation at 700 rcf, washed using PBS, and total RNA was extracted using TRIzol (15596026, Invitrogen) and isopropanol precipitation.

#### Zebrafish growth conditions

Wildtype AB strain zebrafish were maintained in the University of Colorado Anschutz Medical Campus Zebrafish Facility and embryos were staged and maintained according to established protocols (Kimmel et al. 1995). For RNA extraction, 50 embryos at 3 days post fertilization were homogenized in 1 ml of TRIzol (ThermoFisher #15596026). Total RNA was extracted according to the manufacturer’s protocol and eluted in 40 µl nuclease-free water. 10 µg of RNA was used as the input material for deacylation and tRNA isolation. The zebrafish used in this study were approved by the Institution Animal Care and Use Committee under Protocol # 01361.

### Deacylation and tRNA isolation

3-10 µg of total RNA per sample was deacylated in a 50 µL volume with a final concentration of 10 mM Tris-HCl pH 9 for 30 minutes at 30°C. Next, small RNAs including tRNAs were isolated using the Zymo RNA Clean and Concentrator-5 kit (Zymo Research, R1016). Isolation was performed according to the manufacturer-provided instructions to purify molecules 17-200 nucleotides in length, but adding 1.3 volumes instead of the specified 1 volume of 100% ethanol to the flow through containing small RNAs.

### Direct tRNAseq library preparation

To generate matched pairs of ONT RNA002 and RNA004 libraries, 300 ng of deacylated, purified small RNA input was ligated to pre-annealed 5′ and 3′ splint adapters at a 1:1 molar ratio using an in-house preparation of recombinant T4 RNA ligase 2 (homemade preparation,, 0.74 mg/mL). After incubating for 1 hour at room temperature, a 1.8X volume of tRNA purification SPRI beads (BioDynami #40054) was added, and incubated and washed per manufacturer instructions before elution in 9 µL nuclease-free water. This elution product was then ligated to the RTA adapter (Oxford Nanopore Technologies) provided in both the RNA002 and RNA004 sequencing kits using high concentration T4 DNA ligase (NEB, M0202M, 2,000,000 U/mL) at a molar ratio of 3 fold more ligated tRNAs than RTA adapter. After 30 minutes incubation at room temperature, this ligation was also cleaned up with a 1.8X volume of tRNA purification beads and eluted in 25 µL nuclease-free water.

Following the RTA ligation bead cleanup, library yields and ligation efficiency were quality checked via quantification on a Nanodrop spectrometer and running on a TapeStation High Sensitivity D1000 ScreenTape (Agilent). These outputs were used to estimate the molar concentration of each library, which were divided into two samples each containing an equivalent amount of RTA-ligated product (50-100 femtomoles of material) to be input into the final ligation to the RMX (RNA002) or RLA (RNA004) helicase-loaded adapter for 30 minutes at room temperature. These final ligations were cleaned up using a 1.8X volume of Ampure XP SPRI beads (Beckman Coulter), washed with WSB wash buffer (ONT) following the protocol provided with ONT’s SQK-RNA004 and SQK-RNA002 kits.

A complete protocol for the RNA004 tRNA sequencing, including deacylation and tRNA isolation, can be found at https://github.com/rnabioco/tRNA004/.

### Sequencing run conditions

Libraries were run on a combination of MinION instruments and a PromethION P2 Solo instrument connected to a A5000 GPU workstation running MinKNOW software versions 23.07.12 and 23.11.04. SQK-RNA002 libraries were loaded onto R9.4.1 flow cells and sequenced using a custom “short read mode” .toml file based on the simulation parameters detailed in (Lucas et al. 2023), described in more detail at https://rnabioco.github.io/tRNAseq-logistics/. SQK-RNA004 libraries were loaded onto compatible “RNA” flow cells and sequenced using the ≥20 nt setting in MinKNOW for the SQK-RNA004 kit. Common parameters across all runs were the selection of POD5 outputs and disabling of quality filtering to ensure that the lower basecalling quality scores produced at modified nucleotides would not generate additional bias due to read filtering.

More information on sequencing runs and the platform used for each library can be found in Supplemental Table 1.

### Basecalling

Basecalling was performed on all libraries using the open source basecaller Dorado (version 0.7.0) from Oxford Nanopore Technologies (https://github.com/nanoporetech/dorado). RNA002 libraries were basecalled using the “fast” (rna002_70bps_fast@v3) and “high accuracy” (rna002_70bps_hac@v3) models. RNA004 libraries were basecalled using the “fast” (rna004_130bps_fast), “high accuracy” (rna004_130bps_hac) and “super high accuracy” (rna004_130bps_sup) v3.0.1 and v5.0.0 models.

### Run throughput evaluation

To compare sequencing yields across different chemistries, flow cells, and libraries, we used the metric reads per pore per minute (RPPPM), which is generated by extracting the total number of reads per run and the duration of the experiment in minutes from the final line of the `throughput_*.csv` files generated by MinKNOW during each run, as well as the total number of pores found at the point of the initial mux scan from the .html run report. We then divided the total number of reads in the run by the number of pores available, and divided again by the length of the run in minutes to get the number of reads generated per minute for each pore on average.

### tRNA alignment and analysis

tRNA reference files were downloaded from gtRNAdb Data Release 21 (Chan and Lowe 2015) where available. Equivalent fasta files containing high confidence mature tRNA sequences were generated by running tRNAscan-SE 2.0’s general model (Chan et al. 2021) on RefSeq GCF_000189635.1 for *T. thermophila* and NC_000866.4 for enterobacteria phage T4. In all cases, predicted pseudogenes and tRNAs with undetermined (NNN) anticodons were excluded from the reference files. Eukaryotic tRNA fasta references were modified by first appending CCA sequences to each mature tRNA sequence, and then appending and prepending 3′ and 5′ splint adapter sequences, respectively. The *E. coli* reference was treated the same, except CCA addition was omitted as *E. coli* tRNAs contain genomically encoded CCA sequences (Zhu and Deutscher 1987). Finally, a budding yeast mitochondrial reference was built using the experimentally-determined 5′ and 3′ mt-tRNA boundaries described in (Turk et al. 2013), with the addition of CCA and adapter sequences as described above.

Alignments to the tRNA references described above were performed using BWA-MEM version 0.7.16 (Li 2013) with the parameters `bwa mem -C -W 13 -k 6 -T 20 -x ont2d`. Post alignment filtering for full length reads was accomplished using a custom python script (`filter_reads.py`). Coverage per isodecoder was calculated using Bedtools genomecov (Quinlan and Hall 2010) and normalized to counts per million reads before downstream analysis and visualization in Rstudio. Basecalling error statistics were calculated using the custom python script `get_bcerror_freqs.py`. A complete set of analysis scripts, reference sequences, sample data, and Rmarkdown documents to generate plots can be found at the accompanying GitHub repository for this manuscript, <u>github.com/rnabioco/tRNA004/</u>.

## Acknowledgements

This work was supported by the National Institutes of Health (R35 GM119550) and National Science Foundation (Award #2330283). We thank Lydia Heasley for the parent strain and protocol for generating petite yeast, Chad Pearson for *Tetrahymena*, and the members of the Hesselberth lab and RNA Bioscience Initiative community for their constructive feedback throughout this project.

## Data availability

Complete sequencing data (including pod5 and fastq files) are being deposited at the Gene Expression Omnibus; details on the associated BioProject will be included on both the GitHub repository and this manuscript when transfer is complete.

---

**Figure S1.**
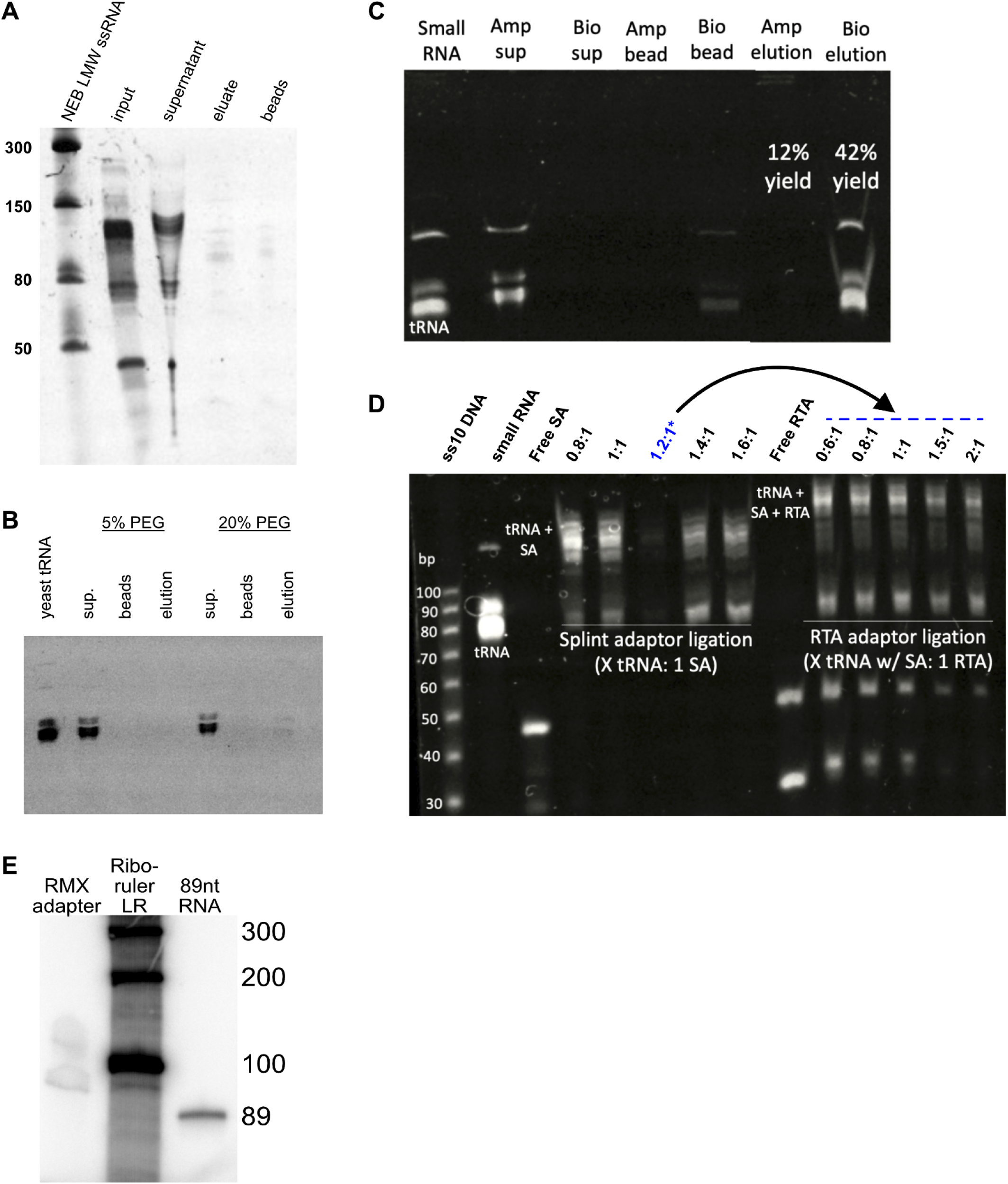
Technical optimizations to improve yields from bead-based cleanups in nanopore tRNA sequencing libraries. (**A**) Ligated and unligated tRNA bind poorly to Ampure XP beads even at a 2.5X bead-to-sample ratio. tRNA ligation reactions to double stranded splint adapters were mixed with Ampure XP beads and incubated per manufacturer instructions. At each step, a 20% volume was removed and retained, including the input, the supernatant after beads were placed on a magnet, the eluate from the beads, and the beads themselves (resuspended in loading dye and loaded directly). All samples were heated to 95°C for 3 minutes in 2X formamide loading dye and then run on an 8% TBU acrylamide denaturing minigel at 250V for 40 minutes before staining with Sybr gold. (**B**) Additional PEG8000 only minimally improves binding of tRNAs to Ampure XP beads. Purified budding yeast tRNA in DEPC H20 was mixed with a 2X volume of Ampure XP beads at a final concentration of 5% or 20% PEG8000 (this final concentration does not include crowding agent already contained within the bead solution). The solution was incubated for 10 minutes at room temperature with rotation, placed on a magnet, washed with 80% EtOH and eluted in DEPC H20. As in **A**, 20% of volume was saved at each step for running on a denaturing gel. (**C**) BioDynami tRNA beads recover substantially more tRNA input than Ampure XP beads. BioDynami (Bio) and Ampure XP (Amp) beads were incubated with purified tRNA at room temperature with rotation, washed with ethanol, and eluted in DEPC H20, with aliquots saved at each step as in the previous panels. (**D**) Optimization of molar ratios for ligation of tRNAs to the double-stranded splint adapter (SA), and the subsequent ligation to the ONT-provided RTA adapter. A 1:1 ratio for ligation 1 and 3:1 ratio for ligation 2 were selected to maximize tRNA capture while minimizing the amount of unligated RTA adapter carried through into the final ligation to a helicase-loaded adapter. Lane annotated in blue was intentionally underloaded on the gel due to use of this material in RTA adapter ligation. (**E**) The helicase-loaded RMX adapter provided in the SQK-RNA002 sequencing kit is approximately 80-100 nt in length. 1 µL of trace RMX adapter was radiolabeled with gamma-ATP using the T4 PNK exchange reaction as detailed in *Molecular Cloning* (Sambrook and Russell 2006), and run on a 6% denaturing acrylamide gel at 250V for 1 hour alongside the radiolabeled ssRNA ladder Riboruler LR and a single stranded 89 nt RNA standard, before drying for 1 hour at 80°C, cooling under vacuum, and overnight exposure of the radioactive gel to a phosphorscreen before imaging on a Typhoon imager (GE). An approximate length of 95 nt was used to estimate the concentration of this reagent along with Qubit high sensitivity DNA assay; based on these experiments, we estimate the helicase-loaded adapter reagents RMX and RLA are provided as 50 nanomolar solutions, and have optimized the molar ratios for the final ligation of RTA-ligated material to these adapters accordingly.

**Figure S2.**
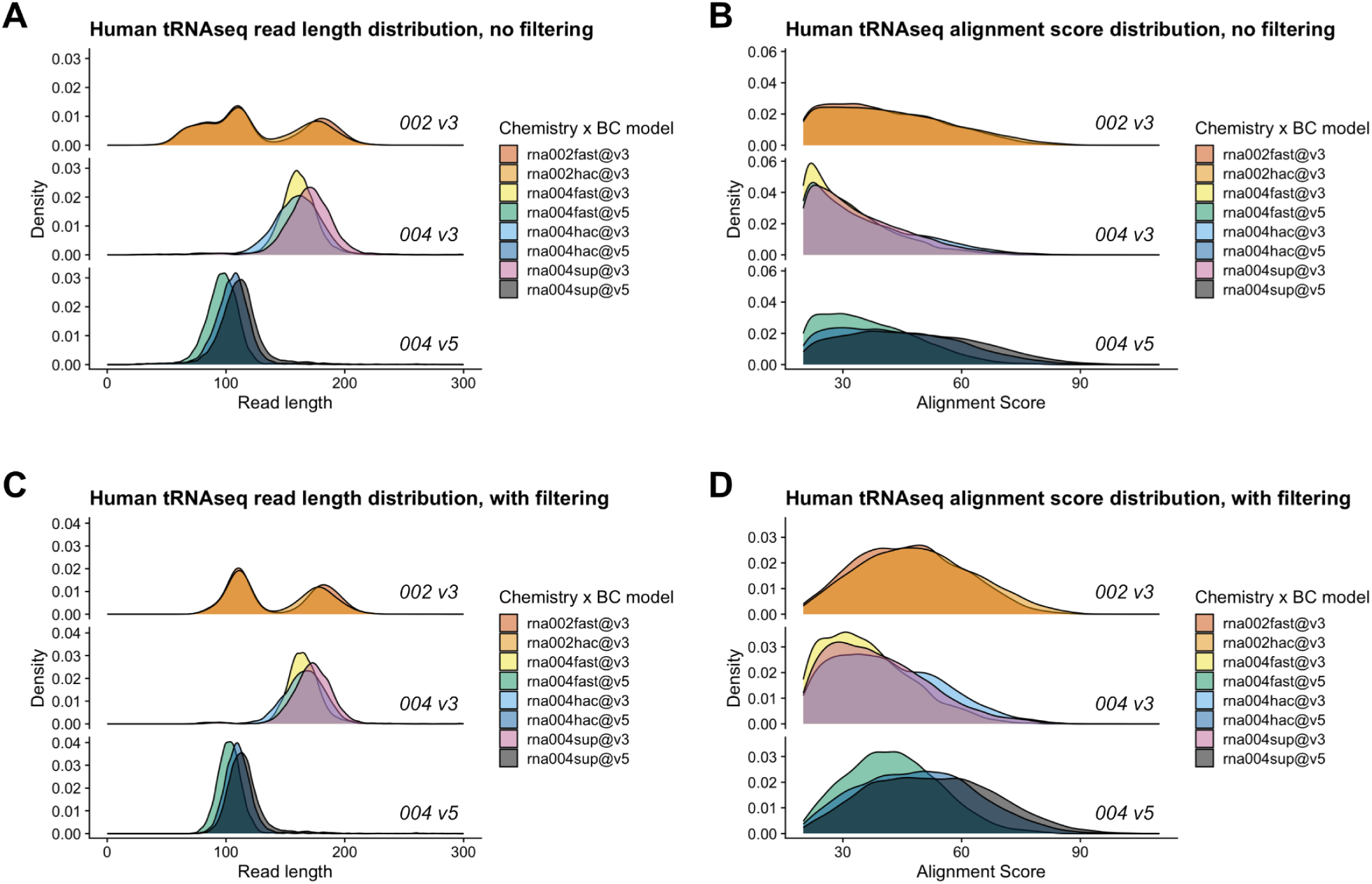
Both tRNA sequencing chemistry and basecalling model selection impact library read length distributions and alignment scores; however, filtering for full length tRNA reads improves alignment scores. Here, human tRNA sequencing libraries subsampled to 10,000 reads provide an illustrative example. (**A**) A comparison of read lengths generated from RNA002 and RNA004 human tRNA sequencing libraries basecalled with Dorado 0.7 using the basecalling models indicated. RNA002 libraries display a multimodal read length distribution, while RNA004 libraries are more uniform, but with longer read lengths produced when *v3* vs. *v5* basecalling models are applied to the same sample. (**B**) tRNA libraries have a broad distribution of alignment scores, but are highest for RNA004 libraries basecalled with the *sup v5* model. (**C**) Post-alignment filtering for full length tRNAs as outlined in Figure 1 reduces RNA002 libraries to a bimodal distribution of read length, and slightly increases the median read length for RNA004 libraries. (**D**) Full length read filtering improves the alignment scores for all combinations of chemistry and basecalling model, with the best performance observed for RNA004 libraries basecalled with the *sup v5* model.

**Figure S3.**
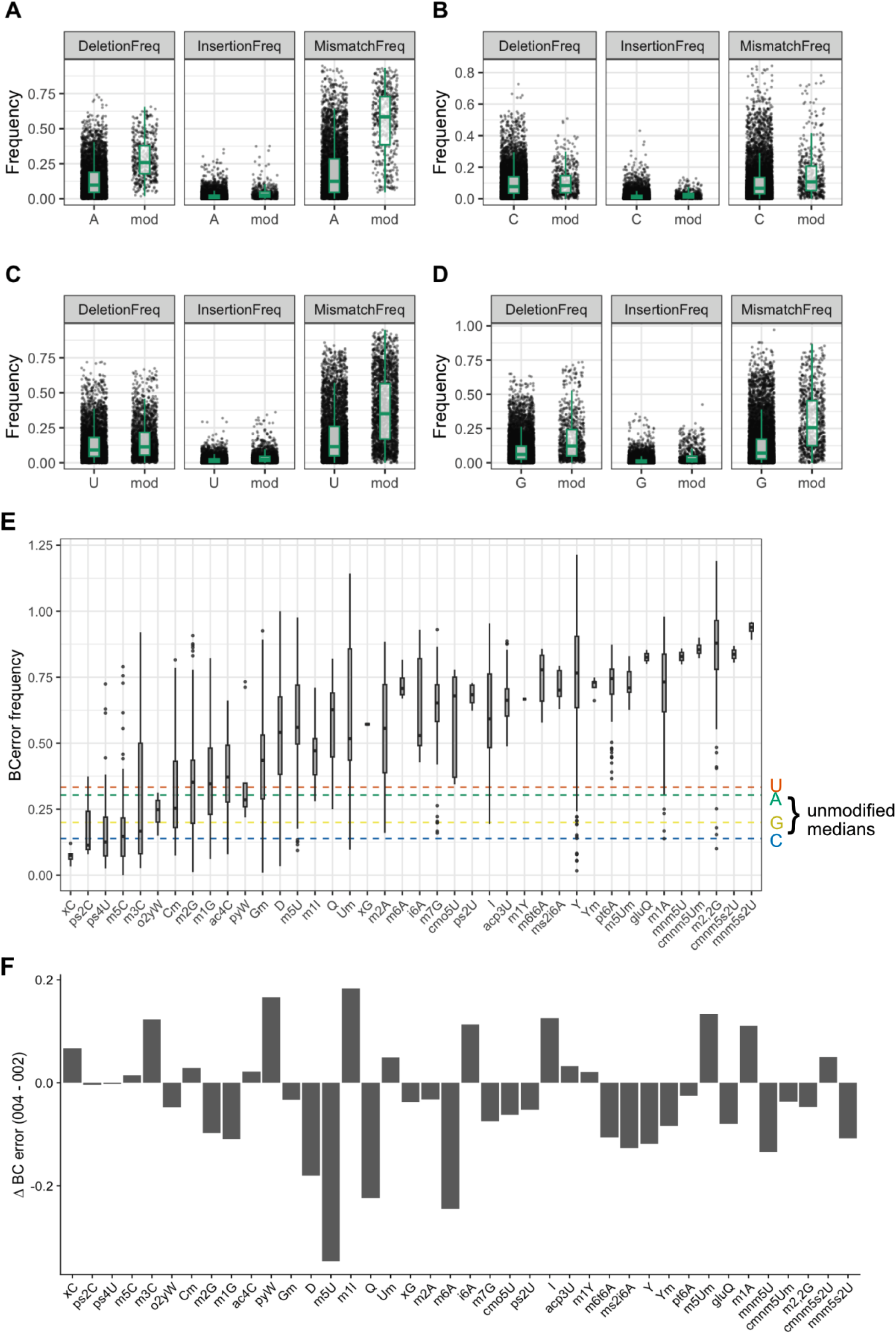
Extended analysis of RNA modification signals. (**A**) Breakdown of basecalling error signal into its component parts (deletion, insertion, and mismatch frequency) for RNA004 libraries basecalled with the DOrado *sup v5* model at modified and unmodified adenosines, (**B**) cytosines, (**C**) uridines, and (**D**) guanosines. (**E**) Distributions of basecalling error frequencies at known modifications on tRNA in libraries sequenced using the first generation RNA002 chemistry and basecalled using the *hac v1* model. As in Figure 2B, modifications are ordered from left to right from lowest to highest median error frequency, with the background levels for unmodified nucleotides in RNA002 sequencing indicated via dashed lines. (**F**) Change in median basecalling error at known sites of tRNA modification with ≥30X coverage in both RNA002 and RNA004 data. Modifications appear in the same order as in panel **E**. Negative values indicate modifications where basecalling error signal is reduced from RNA002 to RNA004 libraries, positive values reflect modifications where error signal has increased with the updated chemistry.

**Figure S4.**
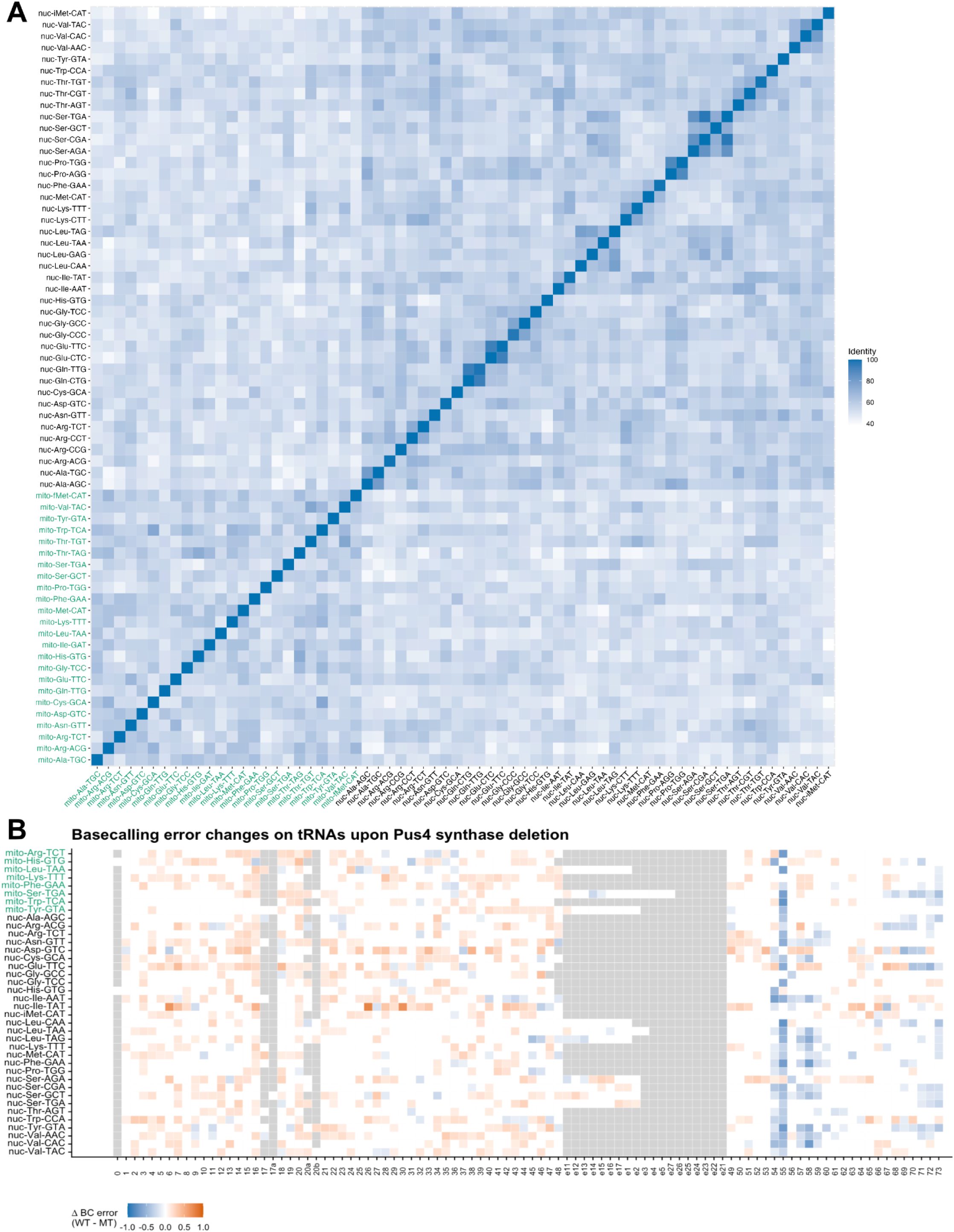
Extended analysis of mitochondrial tRNA signals. (**A**) A pairwise sequence identity matrix for *S. cerevisiae* tRNA sequences encoded in the mitochondrial and nuclear genomes illustrates that mitochondrial and nuclear encoded tRNAs share sequence identity but look most like tRNAs of the same origin. Alignments were generated using the Smith-Waterman algorithm and plotted in R. Mitochondrial tRNAs are indicated in green text. (**B**) Changes in basecalling error between wild-type *S. cerevisiae* and cells with a deletion in the pseudouridine synthase *PUS4* display loss of error signal consistent with loss of modifications at the pseudouridylated position 55 as well as additional modified positions (54, 58) on nuclear tRNAs known to be dependent on nt 55 pseudouridylation. Mitochondrial tRNAs (in green) do not display this same pattern of downstream modification dependencies, but do show negative Δ basecalling error at position 55, consistent with the known nuclear and cytoplasmic distribution of *PUS4*.

**Figure S5.**
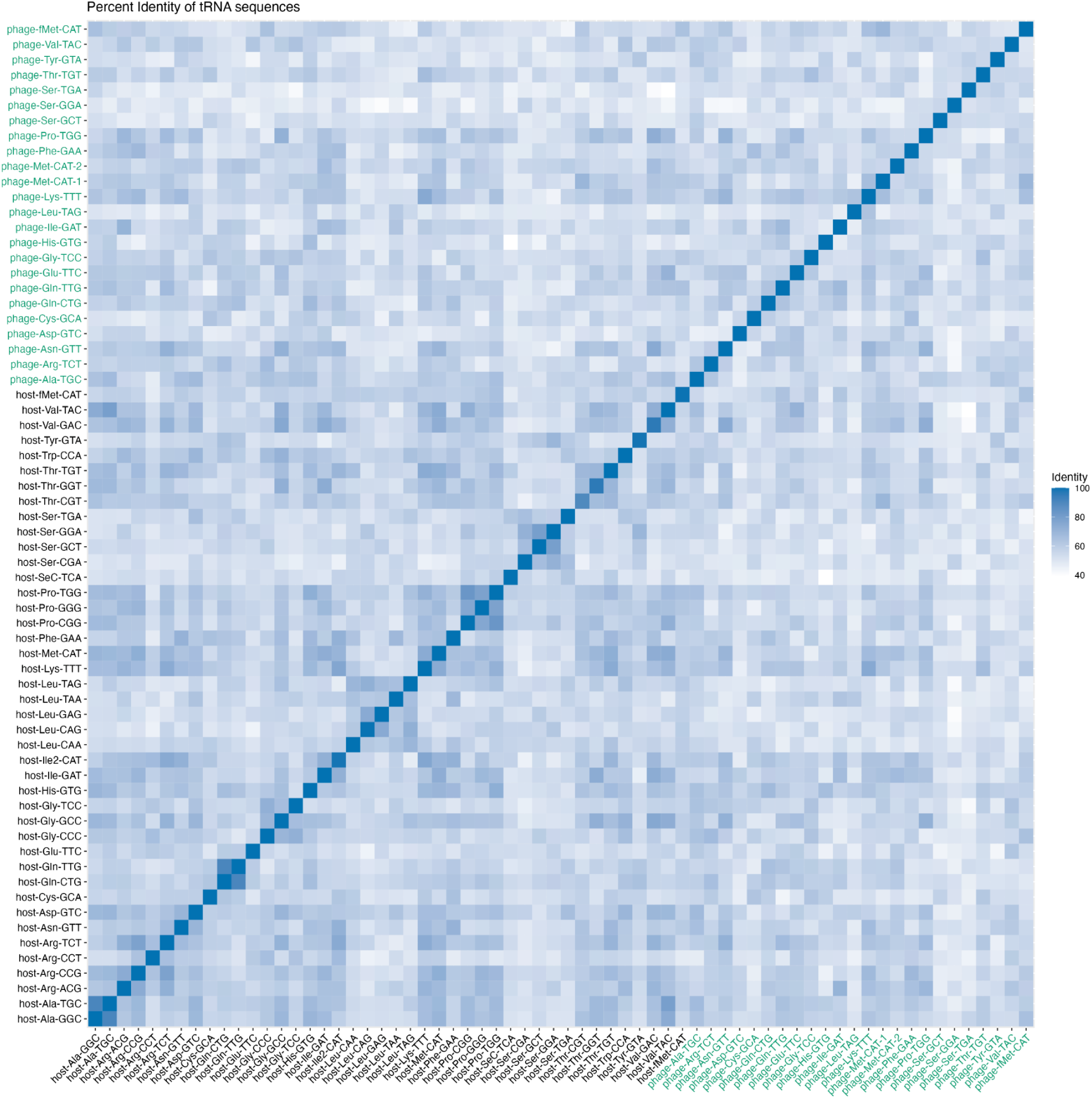
*E. coli* and T4 bacteriophage tRNAs share substantial sequence similarity. As in Figure **S4A**, this plot represents a pairwise sequence identity matrix for tRNAs in the composite reference of T4 phage and *E. coli* tRNA sequences. Smith-Waterman alignments were generated using Parasail and plotted in R. Bacteriophage T4 tRNAs are indicated in green text.

**Table S1.** Sequencing run information.

| Genus | Species | Sample | Chemistry | Platform | Flow Cell ID | Read | Mins | Pores | RPPPM |
| --- | --- | --- | --- | --- | --- | --- | --- | --- | --- |
| Danio | rerio |  | RNA002 | PromethION | PAS44221 | 142321 | 993 | 8832 | 0.02 |
| Danio | rerio |  | RNA004 | MinION | FAX73799 | 199081 | 78 | 1313 | 1.94 |
| Drosophila | melanogaster | S2 | RNA002 | MinION | FAX30455 | 173822 | 114 | 1151 | 1.32 |
| Drosophila | melanogaster | S2 | RNA004 | MinION | FAX73799 | 245444 | 69 | 1052 | 3.38 |
| Escherichia | coli | T4 inf. | RNA004 | PromethION | PAU05281 | 314760 | 53 | 2614 | 2.27 |
| Escherichia | coli |  | RNA002 | PromethION | PAS44628 | 80141 | 157 | 6647 | 0.08 |
| Escherichia | coli |  | RNA004 | MinION | FAX73799 | 253401 | 136 | 504 | 3.7 |
| Homo | sapiens | HS27 | RNA002 | PromethION | PAS29437 | 185971 | 1025 | 7018 | 0.03 |
| Homo | sapiens | HS27 | RNA004 | MinION | FAX71838 | 142888 | 70 | 393 | 5.19 |
| Saccharomyces | cerevisiae | grande | RNA004 | PromethION | PAS98845 | 346152 | 56 | 3602 | 1.72 |
| Saccharomyces | cerevisiae | petite | RNA004 | PromethION | PAQ47538 | 320177 | 56 | 5185 | 1.1 |
| Saccharomyces | cerevisiae | pus1del | RNA004 | PromethION | PAS92267 | 657366 | 69 | 2612 | 3.65 |
| Saccharomyces | cerevisiae | pus2del | RNA004 | PromethION | PAS98920 | 406547 | 69 | 7847 | 0.75 |
| Saccharomyces | cerevisiae | pus4del | RNA002 | PromethION | FAX30455 | 21162 | 999 | 292 | 0.07 |
| Saccharomyces | cerevisiae | pus4del | RNA004 | MinION | FAX71838 | 297027 | 46 | 1066 | 6.06 |
| Saccharomyces | cerevisiae | S288C | RNA002 | PromethION | PAS43478 | 40913 | 31.5 | 10793 | 0.12 |
| Saccharomyces | cerevisiae | S288C | RNA004 | PromethION | PAQ47538 | 204731 | 29 | 7419 | 0.95 |
| Tetrahymena | thermophila |  | RNA002 | MinION | FAX30455 | 557168 | 860 | 1132 | 0.57 |
| Tetrahymena | thermophila |  | RNA004 | MinION | FAX71838 | 296767 | 77 | 1128 | 3.42 |

**Table S2.**
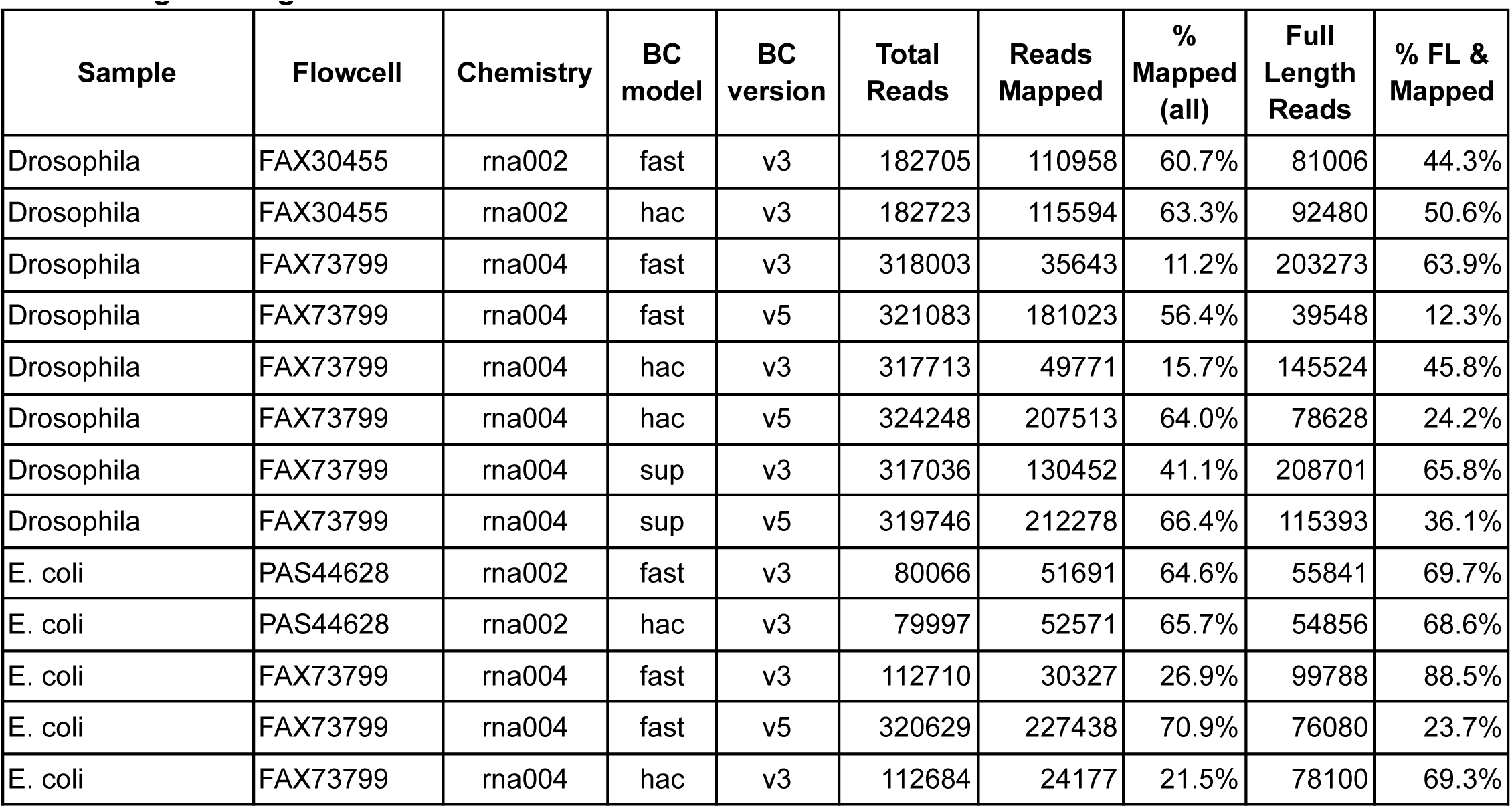
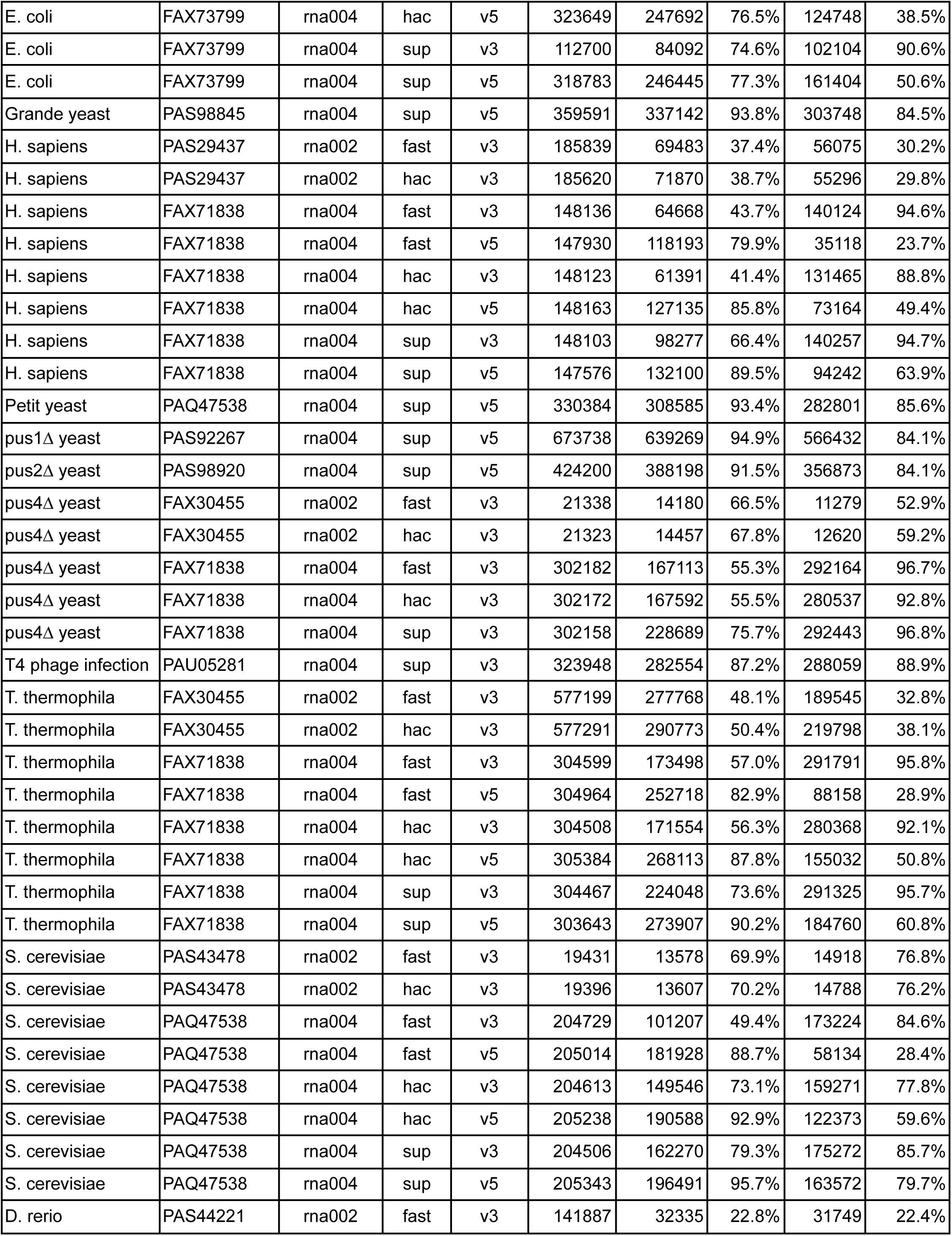
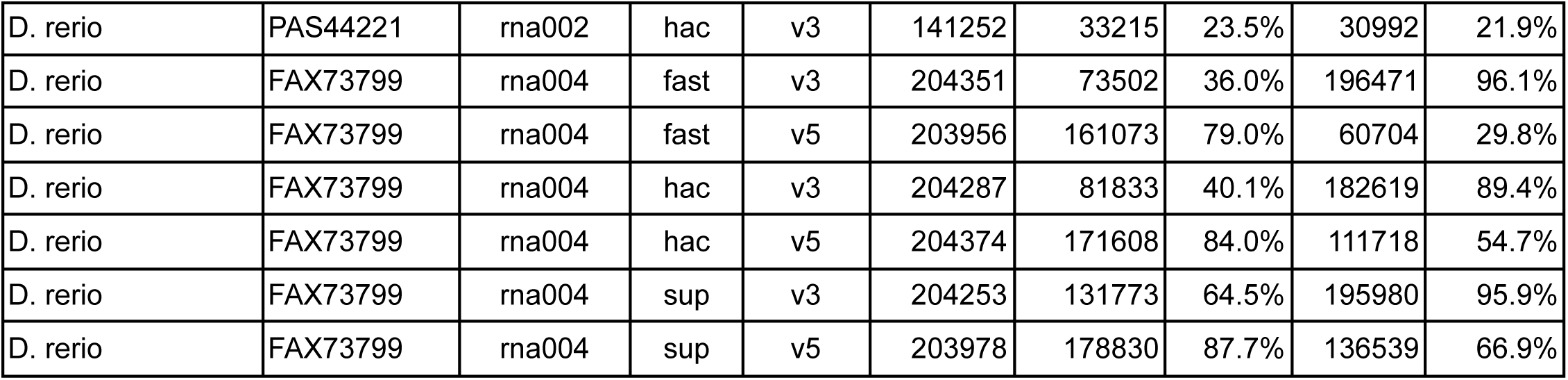
Basecalling and alignment metadata.

**Table S3.**
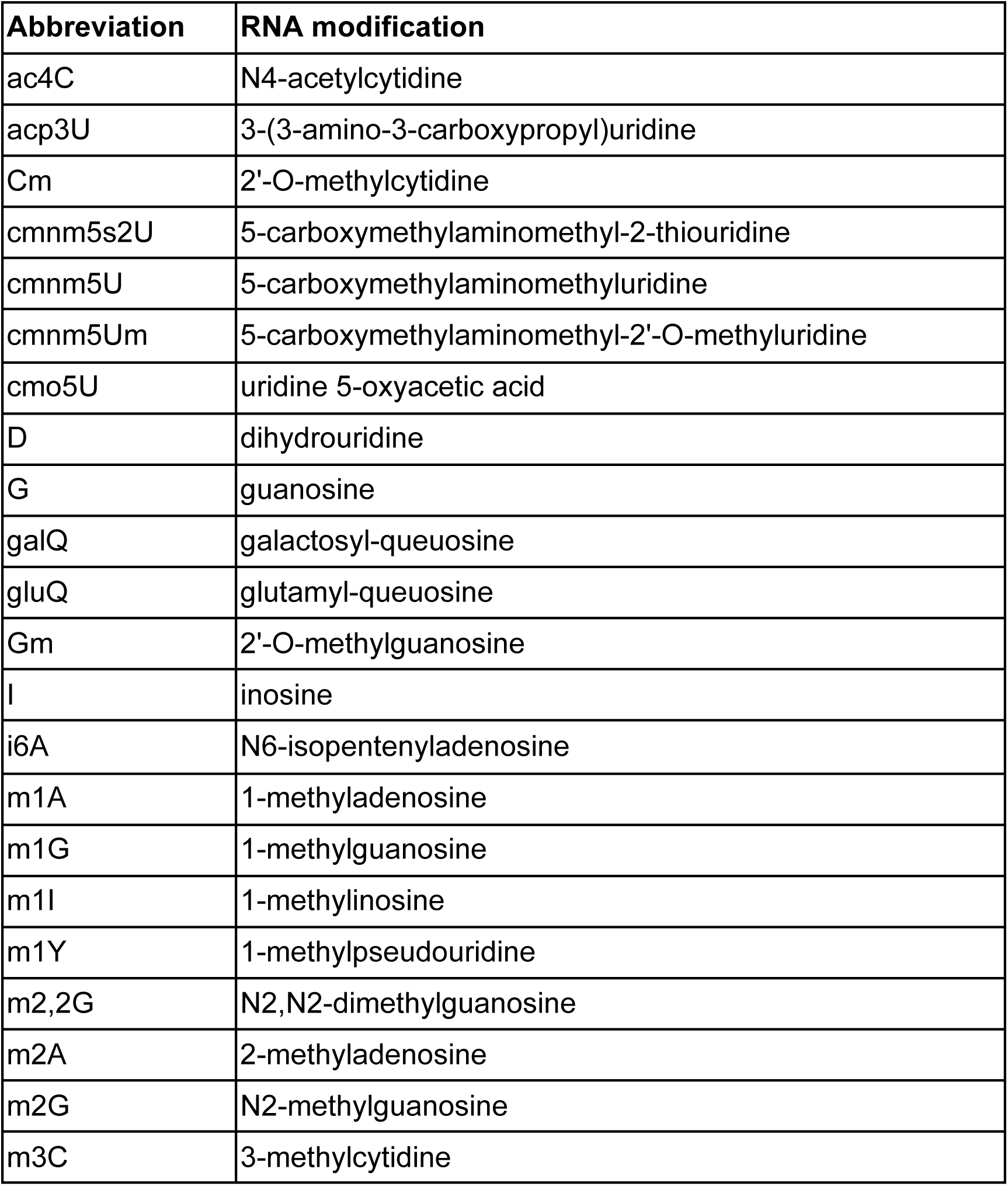
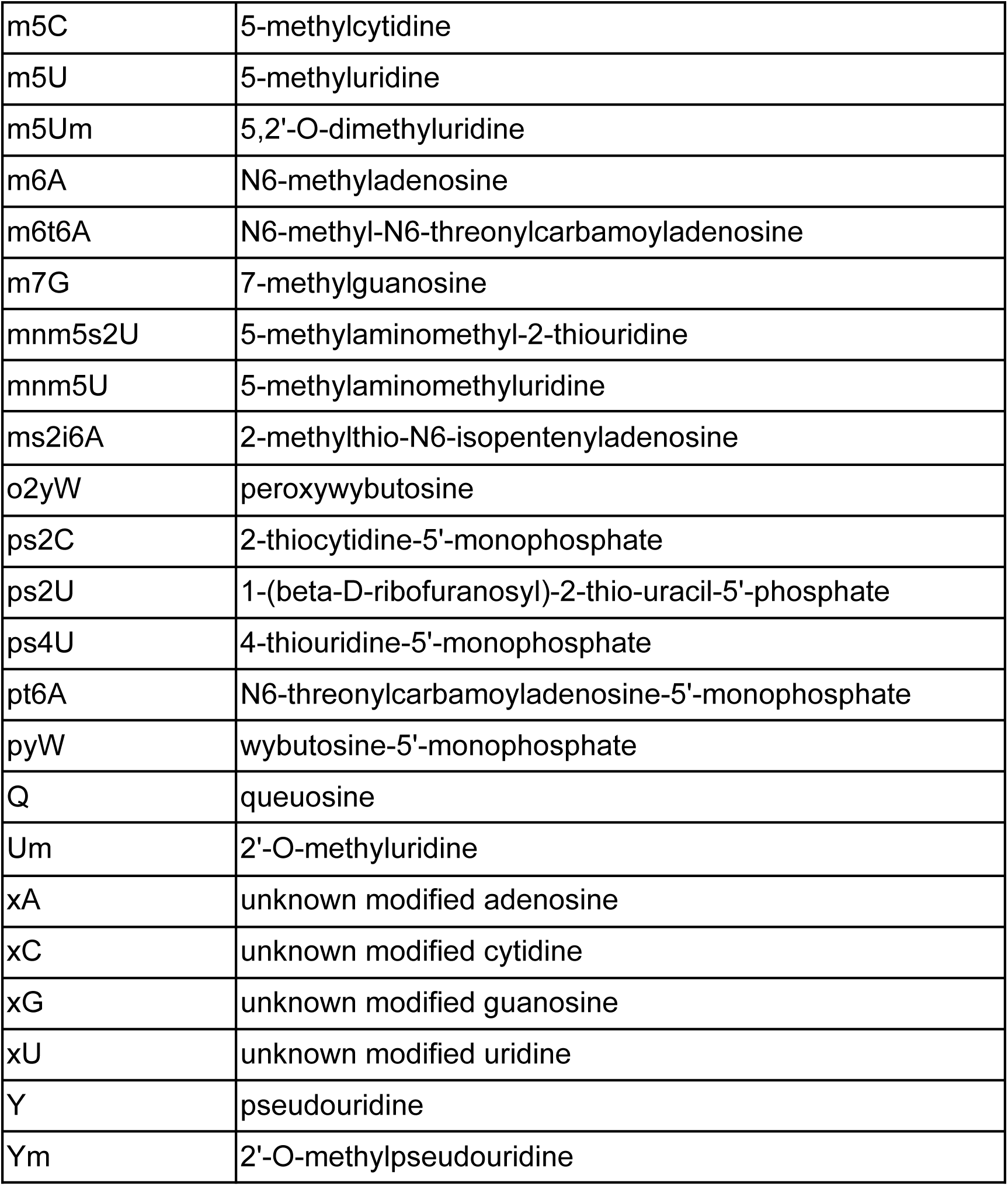
RNA modifications and standardized MODOMICS “short name” abbreviations referred to throughout this text.

